# Regulation of Prefrontal Patterning, Connectivity and Synaptogenesis by Retinoic Acid

**DOI:** 10.1101/2019.12.31.891036

**Authors:** Mikihito Shibata, Kartik Pattabiraman, Belen Lorente-Galdos, David Andrijevic, Xiaojun Xing, Andre M. M. Sousa, Gabriel Santpere, Nenad Sestan

**Affiliations:** Department of Neuroscience, Yale School of Medicine, New Haven, CT 06510, USA; Yale Child Study Center, New Haven, CT 06510; Yale Genome Editing Center, Yale School of Medicine, New Haven, CT 06510, USA; Department of Comparative Medicine, Yale School of Medicine, New Haven, CT 06510, USA; Department of Genetics, Yale School of Medicine, New Haven, CT 06510, USA; Department of Psychiatry, Yale School of Medicine, New Haven, CT 06510, USA; Program in Cellular Neuroscience, Neurodegeneration and Repair, CT 06510, USA; Neurogenomics Group, Research Programme on Biomedical Informatics, Hospital del Mar Medical Research Institute, Department of Experimental and Health Sciences, Universitat Pompeu Fabra, Barcelona, Catalonia, Spain; Kavli Institute for Neuroscience, Yale University, New Haven, CT 06520

**Author notes:** These authors contributed equally to this work (Mikihito Shibata, Kartik Pattabiraman). Correspondence to Nenad Sestan.

## Abstract

The prefrontal cortex (PFC) and its reciprocal connections with the mediodorsal thalamus (MD) are crucial for cognitive flexibility and working memory^1–4^ and are thought to be altered in several disorders such as autism spectrum disorder^5, 6^ and schizophrenia^6–9^. While developmental mechanisms governing regional patterning of the rodent cerebral cortex have been characterized^10–15^, the mechanisms underlying the development of PFC-MD connectivity and the lateral expansion of PFC with distinct granular layer 4 in anthropoid primates^16–23^ have not been elucidated. Here we report increased concentration of retinoic acid (RA), a signaling molecule involved in brain development and function^24, 25^ in the prospective PFC areas of human and macaque, compared to mouse, during mid-fetal development, a crucial period for cortical circuit assembly. In addition, we observed the lateral expansion of RA synthesizing enzyme, ALDH1A3, expression in mid-fetal macaque and human frontal cortex, compared to mouse. Furthermore, we found that enrichment of RA signaling is restricted to the prospective PFC by *CYP26B1*, a gene encoding an RA-catabolizing enzyme upregulated in the mid-fetal motor cortex. Gene deletion in mice revealed that RA signaling through anteriorly upregulated RA receptors, *Rxrg* and *Rarb*, and *Cyp26b1*-dependent catabolism is required for the proper molecular patterning of PFC and motor areas, the expression of the layer 4 marker *RORB*, intra-PFC synaptogenesis, and the development of reciprocal PFC-MD connectivity. Together, these findings reveal a critical role for RA signaling in PFC development and, potentially, its evolutionary expansion.

## Introduction

The proper specialization and expansion of higher-order processing or association areas in the cerebral cortex, specifically PFC, is thought to underlie complex cognitive capabilities. The PFC reaches greatest complexity in anthropoid primates, which have many prefrontal areas that cover the entire anterior two-thirds of the frontal lobe with a well-defined or granular layer 4^16, 17, 19–23^. Mice and rats, the most commonly studied rodents, have fewer prefrontal areas, which are located within the medial agranular (lacking obvious layer 4) frontal cortex that are closest in homology to the primate agranular medial PFC^18, 21^. Given the anatomical and functional divergence of the PFC between primates and rodents, understanding specification of the expanded primate PFC, which likely underlies some of the most distinctly human or primate aspects of cognition, requires analysis of the developing primate cortex. Our previous analyses of the developing human brain revealed that the transcriptomic differences between neocortical areas are transient and most prominent during mid-fetal development^26, 27^, a crucial period for specification and the initial assembly of neocortical neural circuits^28^. Thus, we hypothesized that the molecular processes governing the developmental specification of the human PFC could be revealed by differential regional gene expression analysis of the mid-fetal human neocortex.

### Prefrontal-enriched gene expression gradient in the mid-fetal human neocortex

To identify molecular processes governing the developmental specification of human PFC, we screened for genes that are upregulated in the mid-fetal frontal lobe, using tissue-level RNA-sequencing (RNA-seq) data from BrainSpan and PsychENCODE projects^27^. These data comprise mid-fetal samples from eleven prospective neocortical areas ranging in age from 16 to 22 postconception weeks (PCW), including four PFC areas (medial, mPFC/MFC; orbital, oPFC/OFC; dorso-lateral, dlPFC/DFC; and ventro-lateral, vlPFC/VFC) and the primary motor cortex (M1C). Gene expression in these frontal lobe areas was compared to areas within the parietal (primary sensory cortex, S1C; and inferior parietal cortex, IPC), occipital, (primary visual cortex, V1C) and temporal lobes (primary auditory cortex, A1C; superior temporal cortex, STC; and inferior temporal cortex, ITC) (Fig. 1a; Extended Data Fig. 1a). We identified 190 protein-coding genes that are specifically upregulated in at least one area within a lobe in comparison with areas from other lobes: 125 in the frontal lobe, 46 in the temporal lobe, 17 in the parietal lobe, and 2 in the occipital lobe (Extended Data Fig. 1a,b). The expression profiles of these 190 differentially expressed genes were able to differentiate the four brain lobes and most regions within them, as observed by principal component analysis (PCA). Moreover, the first principal component (PC1), the one accounting for the highest variability present in the data, corresponded to the anterior-posterior axis of the neocortex with genes specifically upregulated in the frontal lobe mainly negatively correlated with PC1 while genes upregulated in the other lobes are positively correlated with PC1 (Fig. 1d). Interestingly, Gene Ontology (GO) enrichment analysis of the 125 genes upregulated in the mid-fetal frontal lobe identified a significant enrichment of genes associated with categories such as response to retinoic acid (RA), synapse development and axon development/guidance (Fig. 1c; **Supplementary Table 1**), suggesting that these genes may play a role in frontal lobe patterning and circuit development. Accordingly, multiple frontal lobe upregulated genes associated with RA signaling, such as *CBLN2*, *RXRG*, *CBLN1*, *NEUROD1*, *NEUROG2*, *CDH8*, *CYP26A1,* and *MEIS2*^24, 25, 29–31^ (**Supplementary Table 2**; see also this and our accompanying study^31^), were amongst those with the highest impact on PC1 (Fig. 1d).

**Figure 1.**
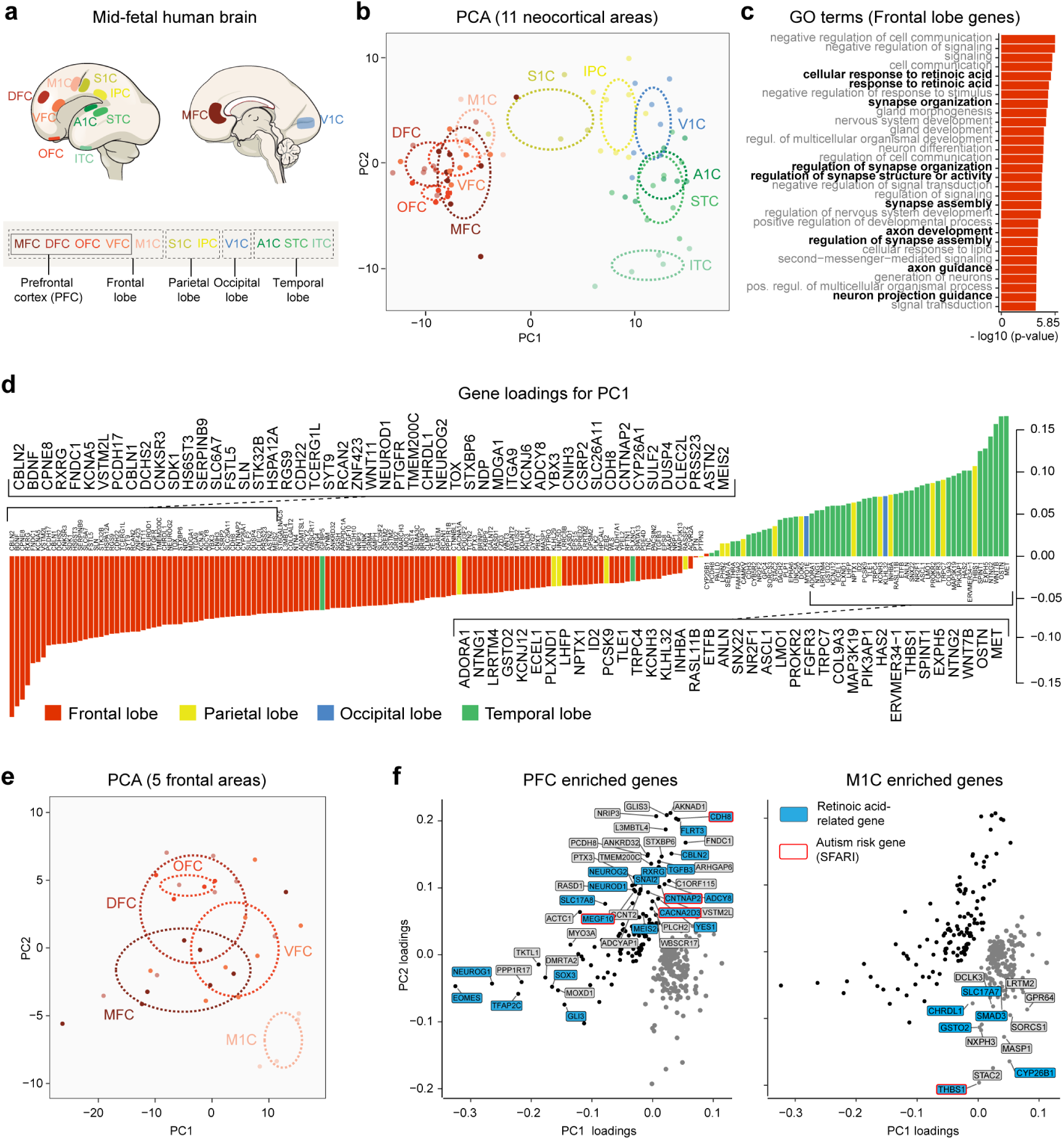
Retinoic acid-related genes are upregulated in human mid-fetal PFC. **a,** Diagram of the 11 human mid-fetal neocortical areas from the four brain lobes used for analysis. **b**, First two principal components based on the expression of 190 protein-coding genes that are specifically upregulated in one of the four brain lobes. Each color represents a neocortical area. Ellipses are centered on the mean of the points of a given area and the size of the axes corresponds to their standard deviation on each component. **c**, GO terms associated with the genes specifically upregulated in the frontal lobe and their nominal P value. **d**, Gene loadings of the first principal component. Colors represent the brain lobe where the gene was found to be specifically upregulated. **e**, First two principal components based on the expression of 306 genes differentially expressed between PFC and M1C. **f,** Loadings of the first two principal components with labeled genes being upregulated in PFC (black dots) and M1C (gray dots). Genes highlighted in blue are associated with RA signaling, and genes with red outline have ASD risk alleles.

As the frontal lobe consists of both prefrontal association and motor regions, we checked whether RA-related genes were upregulated in the PFC as compared to the prospective M1C. Analyzing only the RNA-seq samples encompassing the five areas of the mid-fetal human frontal lobe from Fig.1a^27^, we identified 116 protein-coding genes upregulated in at least one of the four prospective PFC areas and 190 genes upregulated in the prospective M1C. Together, these 306 differentially expressed genes distinguished M1C from PFC as shown by PCA (Fig. 1e). RA-related genes, such as *CBLN2, RXRG*, *CDH8, MEIS2*, and *RBP1* were among the genes upregulated in the PFC compared to the M1C, while the RA degrading enzyme *CYP26B1* was upregulated in the M1C compared to the PFC (Fig. 1f; **Supplementary Table 2**), consistent with our previous microarray-based findings in the human mid-fetal cortex^32^. Upregulated expression of *RXRG* and *CBLN2* in prospective mid-fetal PFC were confirmed by *in situ* hybridization in human as well as in rhesus macaque, the most commonly studied non-human anthropoid primate, and mouse, at developmental ages equivalent to the human mid-fetal period^28^ (Extended Data Fig. 2c; see also our accompanying study^31^). We also identified multiple autism spectrum disorder (ASD) risk genes, such as *CACNA2D3*, *CDH8*, *CNTNAP2*, and *MEGF10* (Group 1-3 from SFARI Gene database, gene.sfari.org), upregulated in the mid-fetal PFC (Fig. 1f; **Supplementary Table 2**).

**Figure 2.**
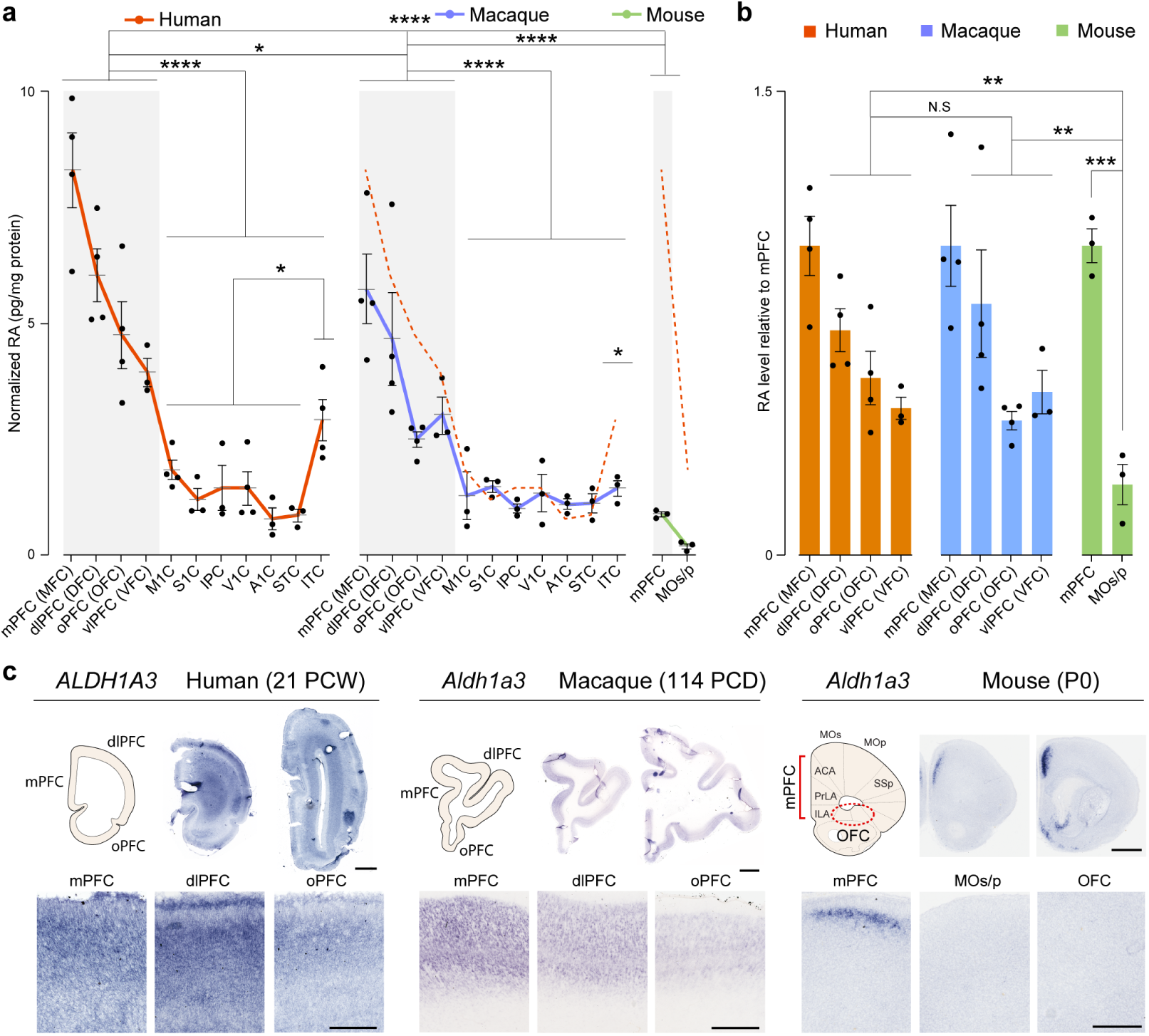
Lateral expansion of the retinoic acid synthesizing enzyme, ALDH1A3 in primate mid-fetal frontal lobe. **a**, Quantification of RA concentration by ELISA from 11 cortical areas from all four neocortical lobes of the human (16, 18, 19, 21 PCW) and macaque (Four 110 PCD brains), and mPFC and MOs from mouse (Four P1 brains) (N = 3-4 for each areal sample). Details about dissection of postmortem brain subregion samples are described in Methods. RA amount was normalized by the total protein in the lysate. Two-tailed unpaired t-test: ****P < 5e-5, *P *<* 0.05; N = 3-4 per condition; Errors bars: S.E.M. Dashed red line in macaque and mouse graphs represent human RA concentration. **b**, Relative RA level in mPFC, dlPFC, vlPFC compared to mPFC (human, macaque) or mPFC, MOs compared to mPFC (mouse). Two-tailed ratio paired t-test: ***P = 0.0004, ** P < 0.005, N = 3-4; Errors bars: S.E.M. **c**, Schema of the PFC and representative images of cortical *ALDH1A3* expression in coronal sections of human (21 PCW), macaque (114 PCD), and mouse (P0) brains using *in situ* hybridization. Expression in human and macaque mPFC, dlPFC, and oPFC, and mouse mPFC comprised of infralimbic (ILA), prelimbic (PrLA), and anterior cingulate area (ACA), MOs/p, and OFC are shown at higher magnification. Note that *ALDH1A3/Aldh1a3* expression is expanded to all regions of the anterior part of the mid-fetal human and macaque frontal cortex (PFC), while it is restricted to the medial frontal cortex (mPFC) in mouse. Scale bars: 200 μm (mouse); 2 mm (human and macaque); 100 μm (mouse, lower panel); 500 μm (human and macaque, lower panel). N = 2 for human and macaque; N = 3 for mouse.

### Prefrontal-enriched retinoic acid concentration gradient in mid-fetal primates

RA is a diffusible biologically active derivative of vitamin A previously implicated in neurogenesis, differentiation and synaptic function of cortical neurons^24, 25, 29, 30, 33–38^. RA is also known to exert a trophic influence on nascent neurons to promote axon and dendritic outgrowth ^33, 35, 39, 40^ as well as synaptogenesis^35, 41^. Moreover, alterations in RA signaling have been implicated in the pathophysiology of ASD^42–44^ and schizophrenia^45–48^. Given the enrichment of RA-related genes among those upregulated in the mid-fetal human PFC, we assessed whether RA concentration is increased in the PFC, using an RA-specific enzyme-linked immunosorbent assay (ELISA) in two anthropoid primates with laterally expanded PFC, human and macaque, and mouse, a rodent with fewer, medially located, PFC areas. Quantification of RA concentration in the same eleven mid-fetal neocortical areas analyzed for differential gene expression revealed a prominent PFC-enriched anterior-posterior gradient of RA concentration in postmortem human (16,18,18,19 PCW; N = 3-4 brains per areal measurement) and macaque (four brains at 110 postconceptional days (PCD); N = 3-4 brains per areal measurement) neocortex, with mPFC exhibiting the highest concentration within each species (Fig. 2a). Overall, RA concentrations were much higher in prospective PFC areas (mPFC, dlPFC, oPFC and vlPFC) compared to more posterior areas (M1C, S1C, IPC, V1C, A1C, STC, and ITC; Two-tailed unpaired t-test: P = 2e-6; N = 15 in PFC areas, 21 in posterior areas). Complementary analysis of mouse neonatal cortex revealed higher RA concentrations in prospective mPFC compared to the adjacent secondary and primary motor areas (P1; N = 3 brains; Fig. 2a).

Comparison across the three species at approximately equivalent developmental age, identified higher concentrations of RA in both the human mPFC and all four PFC areas overall as compared to the other two species (Two-tailed unpaired t-test: Human all PFC vs. macaque all PFC: 1.45 fold change, P = 0.02; human mPFC vs macaque mPFC: 1.45 fold change, P = 0.03; human all PFC vs mouse mPFC: FC 6.48, P = 3e-8; human mPFC vs mouse mPFC: 9.34 fold change, P = 0.0005; Fig. 2a), as well as in macaque compared to mouse (Unpaired t-test: Macaque all PFC vs mouse mPFC: 4.48 fold change, P = 2e-6; macaque mPFC vs mouse mPFC, 6.45 fold change, P = 0.003; Fig. 2a). Interestingly, ITC, an association area within the temporal lobe thought to exhibit unique features and connectivity in humans ^23, 49^, had a higher RA concentration among non-frontal areas in humans but not in macaque (Unpaired t-test: P = 0.01; Fig. 2a).

### Lateral expansion of the mid-fetal expression of retinoic acid synthesizing enzyme in primates

Next, we assessed how RA concentration differs across the medial-lateral axis of the mid-fetal frontal lobe in human and macaque compared to the mouse frontal cortex at the equivalent developmental age. In mice, we detected increased RA concentration in the medial frontal cortex, which comprises PFC areas (mPFC), compared to adjacent dorso-lateral regions corresponding to prospective secondary and primary motor areas (Fig. 2a). The ratio of the concentration of RA between the dorsal and medial PFC/frontal cortex was elevated in human and macaque compared to mouse, with no significant difference between human and macaque (Two tailed ratio paired t-test: human vs. mouse, P = 0.003; macaque vs. mouse, P = 0.003; human vs. macaque, NS; Fig. 2b), indicating that RA signaling is expanded laterally in the human and macaque mid-fetal frontal cortex, compared to mouse.

The observed enrichment of RA in the PFC led us to examine the mid-fetal expression of a key RA synthesizing enzyme, ALDH1A3/RALDH*3* ^24, 25, 53^ in human, macaque and mouse. The mPFC enrichment of RA concentration in all three species (Fig. 2a, b), closely overlaps with previously reported expression of ALDH1A3 in mice^54^. We confirmed this in the neonatal mouse frontal cortex (N = 3 brains) and found *Aldh1a3* expression predominantly in the upper layers of the dorsal portions of the mPFC and the cingulate cortex (Fig. 2c). However, unlike the medially restricted expression of *Aldh1a3* in mice, we identified low but spatially broader expression of human *ALDH1A3* (21 and 22 PCW; N = 2 brains) throughout the cortical plate, underlying subplate zone and white matter in all regions of the anterior part of the mid-fetal frontal lobe, which correspond to the prospective areas of the medial, orbital and lateral PFC (Fig. 2c). A similar pattern of regional expression was observed throughout the cortical plate of the macaque cortex (114 and 140 PCD; N = 2 brains), but with less noticeable expression in the subplate zone and white matter, compared to human PFC (Fig. 2c). This finding indicates that the expression domain of *ALDH1A3* has expanded dorso-laterally and ventro-laterally in human and macaque mid-fetal frontal lobe, similar to lateral extension of granular PFC in anthropoid primates^16, 20^.

### Developmental prefrontal retinoic acid signaling is mediated by RXRG and RARB

Given the mid-fetal anterior cortical upregulation of both RA and ALDH1A3, we further assessed the expression of RA-dependent receptors and RA-responsive downstream genes in the human, macaque and mouse cortex. In addition to *Rxrg*, which is upregulated in the human mid-fetal PFC (Fig. 1f and Johnson et al. ^32^), we also identified an upregulation of *Rarb* in the P0 mouse anterior cortex by quantitative PCR (Extended Data Fig. 2a,b). *In situ* hybridization revealed that both *RXRG* and *RARB* orthologs are more highly expressed in the striatum as compared to the cerebral cortex. However, within the mid-fetal human (21, 22 PCW; N = 2 brains) and macaque (114, 140 PCD; N = 2 brains) cortex, or neonatal mouse cortex (P0; N = 3 brains), the two genes exhibited a high anterior to low posterior gradient of expression (Extended Data Fig. 2c).

The RXRG/RARB heterodimer has been previously shown to mediate RA signaling in the adult mouse cerebral cortex and striatum, and be required for learning, locomotion and dopamine signaling^29, 30^. To assess whether RXRG and RARB are required for RA signaling activity in the developing mouse frontal cortex, we generated constitutive *Rxrg* and *Rarb* double knockout (dKO) mice (Extended Data Fig. 2d), which, consistent with previous findings^29, 30^, are viable. To assess RA signaling in these dKO mice, we crossed these mice with the *RARE-lacZ* reporter line^52^ where *lacZ* is under the transcriptional control of a RA response element (RARE). Using control *RARE*-*lacZ* mice we first confirmed previously described enrichment of RA signaling in the dorsal portion of medial frontal and limbic areas in neonatal mouse^51^ (Fig. 3a; Extended Data Fig. 3a). We then identified a reduction of RA signaling in the neonatal dKO mPFC (Two-tailed Student’s t-test: WT vs. dKO, P *=* 1e-6 in section 1; P *=* 0.0003 in section 2 in Fig. 3a, Extended Data Fig. 3a). In addition, we found a less extensive reduction of RA signaling in the anterior cingulate area (ACA) and retrosplenial area at P0 (Extended Data Fig. 3a). We also observed reduction of RA signaling in the hippocampus and outer shell of the striatum of dKO mice (Extended Data Fig. 3a). There was no reduction of signaling in the thalamus, where expression was restricted to ventroposterior nuclei (Extended Data Fig. 3a). Furthermore, expression of genes upregulated in the human mid-fetal and mouse neonatal frontal lobe and known to be regulated by RA^31, 54^, including *Cbln2* and *Meis2* (Fig. 1e), were reduced in the dKO mice (Extended Data Fig. 3b). These findings indicate that RXRG and RARB are required for RA signaling in the neonatal mouse mPFC.

**Figure 3.**
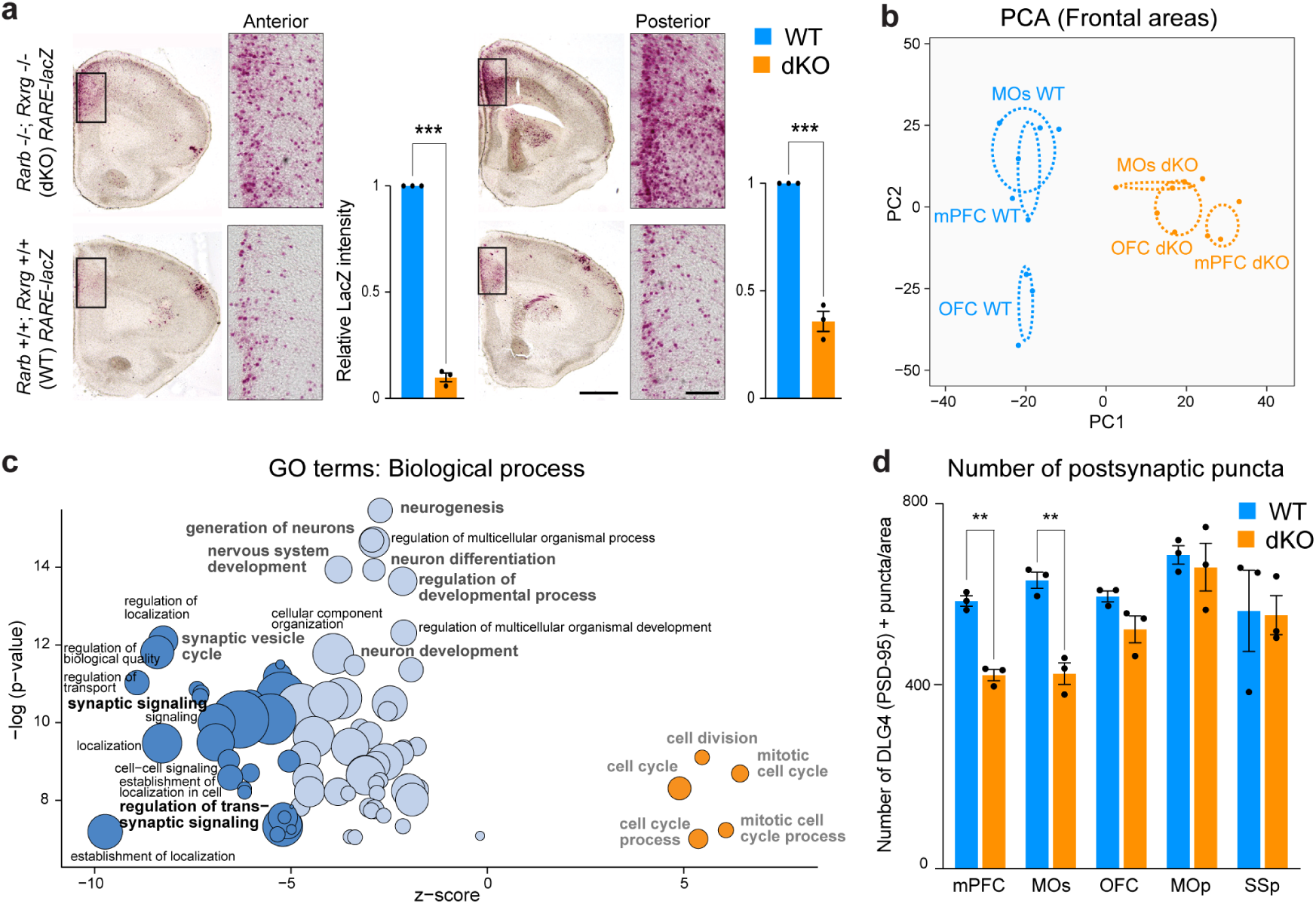
Reduced RA signaling in the PFC of mice lacking RXRG and RARB leads to downregulation of genes involved in synapse and axon development. **a**, β-Galactosidase histochemical staining in the mouse PFC from *Rxrg*+/+; *Rarb*+/+; (WT); *RARE-lacZ* (Blue) and *Rxrg*-/-; *Rarb*-/-(dKO); *RARE-lacZ* (orange) brains at P0. In the dKO brains, LacZ signal is reduced in the mPFC (inset) (N = 3 per genotype). Two-tailed Student’s t-test: WT vs. dKO: ***P *<* 0.0005; Errors bars: S.E.M.; Scale bars: 200 μm; 50 μm (inset). **b,** First two principal components calculated from the expression of 4,768 protein-coding genes that are differentially expressed between WT and dKO mice in at least one of the three frontal cortex areas (mPFC, MOs, and OFC). **c,** GO terms associated with the 4,768 DEx genes showing their z-score and nominal P values. Z-score represents the proportion of upregulated versus downregulated genes in the dKO compared to WT in the list of DEx genes associated to each GO term (i.e. z-score = (#up - #down) / sqrt (#all DEx associated to the GO term)). Dark blue: z-score < −5; light blue: z-score (−5,0]; orange: z-score >0. Size of the bubbles are proportional to the total number of DEx genes associated to the given GO term. **d**, Quantification of excitatory synapses marked by PSD-95/DLG4 in the mPFC, MOs, OFC, MOp, and SSp regions from P0 WT and dKO mice brains. Two-tailed Student’s t-test: **P < 0.005; Errors bars; S.E.M.; N = 3 per genotype.

### RXRG and RARB regulate prefrontal synaptogenesis and axon development

To understand the functional significance of RA signaling through the RXRG/RARB heterodimer in the developing cortex, we performed RNA-seq analysis of different regions/areas of the P0 mouse frontal cortex (i.e., mPFC; secondary motor cortex (MOs) and the adjacent parts of the primary motor cortex (MOp); and OFC, as indicated in Fig. 2c) microdissected from postmortem *Rarb* and *Rxrg* dKO and WT littermate control brains. We identified 4768 differentially expressed (DEx) protein-coding genes between the two genotypes in at least one of the areas (Extended Data Fig. 4a). The highest number of DEx genes was found in the mPFC, as expected based on the medial frontal enrichment of RA signaling in P0 mice (mPFC: 2630 genes; OFC: 2095 genes; MOs/MOp: 1240 genes; Extended Data Fig. 4a; **Supplementary Table 3**). PCA based on the expression of these DEx genes separated the WT and dKO along PC1, with the mPFC showing greatest distance between the WT and the dKO, further supporting the notion that the mPFC was most affected by the reduction of RA signaling (Fig. 3b, Extended Data Fig. 4b).

**Figure 4.**
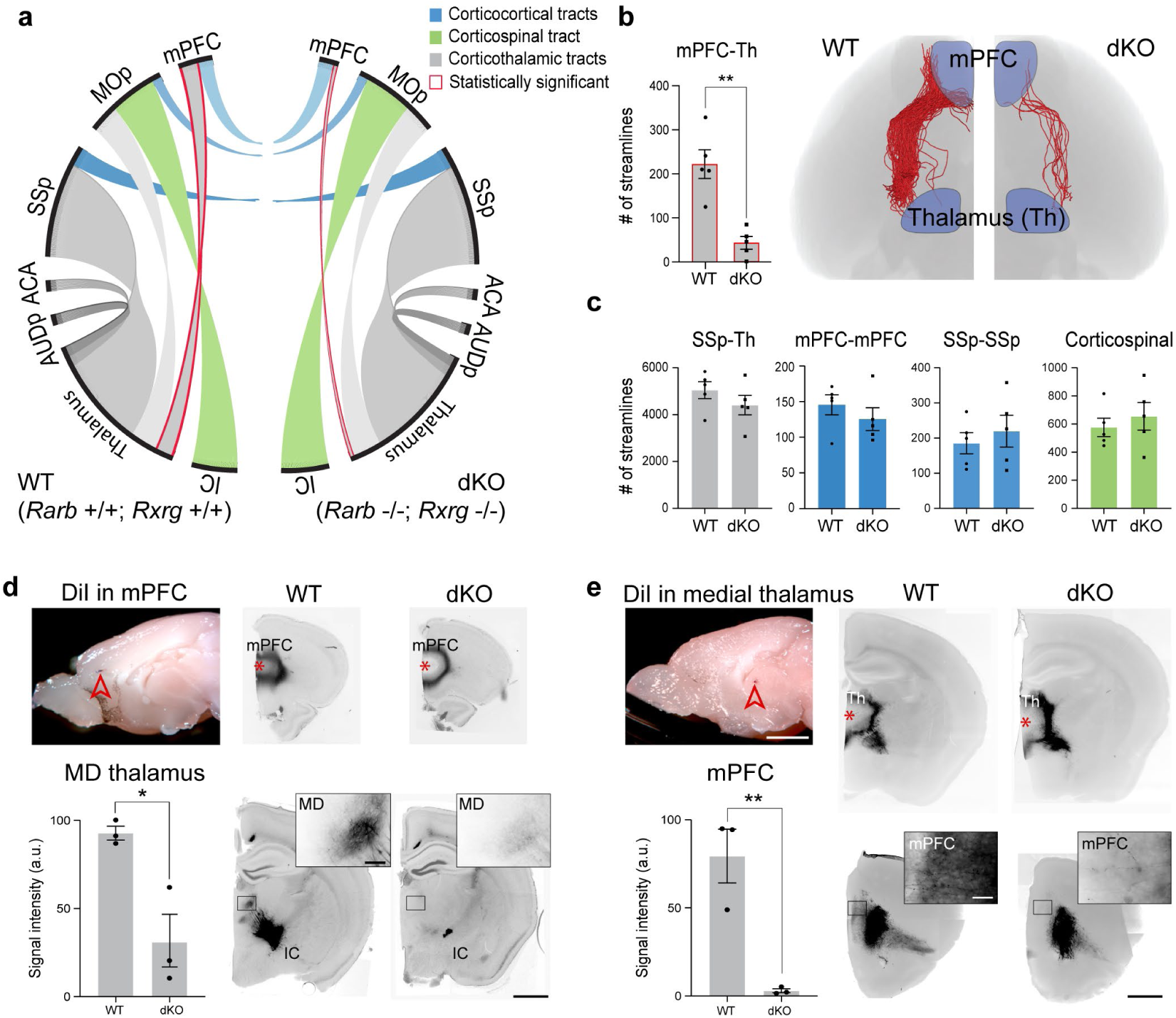
Loss of mPFC-MD thalamic connections in mice lacking *Rarb and Rxrg*. **a,** Representation of number of streamlines (NOS) generated as a connectivity measurement between the cortical areas, thalamus and internal capsule (IC) at P5 using DTI. In dKO brains, there is a significant decrease in NOS generated between the thalamus and mPFC. NOS between other regions are not significantly different between dKO and WT brains (N = 5 per genotype). **b,** Visualization and quantification of streamlines between mPFC and thalamus in dKO and the WT brains. Paired t-test: **P < 0.005; N = 5 per genotype. **c**, Quantification of select corticothalamic, corticocortical and corticospinal streamlines. Paired t-test: NS; Errors bars: S.E.M; N = 5. **d**, **e**, DiI placement in the mPFC and medial thalamus in WT and dKO brain at P21. In dKO brains, the labelled processes detected in the mediodorsal (MD) thalamus (inset), and mPFC (inset) are significantly reduced, indicating the reduction of reciprocal thalamocortical and corticothalamic projections. Two-tailed Student’s t-test: *P = 0.009, **P = 0.003; N = 3 per genotype; Errors bars: S.E.M.; Scale bars: 1 mm; 100 μm (inset).

The GO enrichment analysis revealed that terms associated with genes that are overexpressed in the WT frontal cortex compared to the dKO frontal cortex were highly related to the process of synaptogenesis and cellular components related to synapses and axons, whereas the ones overexpressed in the dKO are related to the regulation of the cell cycle (Fig. 3c; Extended Data Fig. 4c,d). In addition, reflecting the enrichment of RA signaling in the medial frontal cortex (Fig. 3c), when analyzing DEx genes in different regions/areas of the frontal cortex, only genes overexpressed exclusively in the mPFC were associated with the cellular components, axons and synapses (Extended Data Fig. 4c,d). Of note, the majority of DEx genes with GO terms related to axon and synapse development showed the presence of RA receptor binding sites in their associated regulatory elements (promoters, putative enhancers) with genes with reduced expression in the dKO more likely to have at least one RA receptor binding sites (Extended Data Fig. 5). We extended this analysis to the human mid-fetal frontal lobe upregulated genes related to axons and synapses, and identified the presence of RA receptor binding sites in their associated regulatory elements (Extended Data Fig. 5). However, binding site enrichment analyses using two independent methods did not reveal over-representation of RA receptor binding sites compared to random sequences or transcriptional start site’s flanking regions of human genes for either set of genes (Extended Data Fig. 5). Furthermore, several of the genes downregulated in the dKO related to axon guidance (such as *Chl1, Epha8, Efnb3, Epha4, Ephb6, Ntf3, Robo1, Nr4a2, Plxnc1,* and *Plxnd1*) and synapse development (such as *Cbln2*, *Cdh8*, and *Wnt7b*) displayed an anterior-enrichment in WT neonatal mouse cortex (Extended Data Fig. 6a; **Supplementary Table 4**; see also our accompanying study^31^), similar to gradient of *Rxrg* and *Rarb* (Extended Data Fig. 2c).

**Figure 5.**
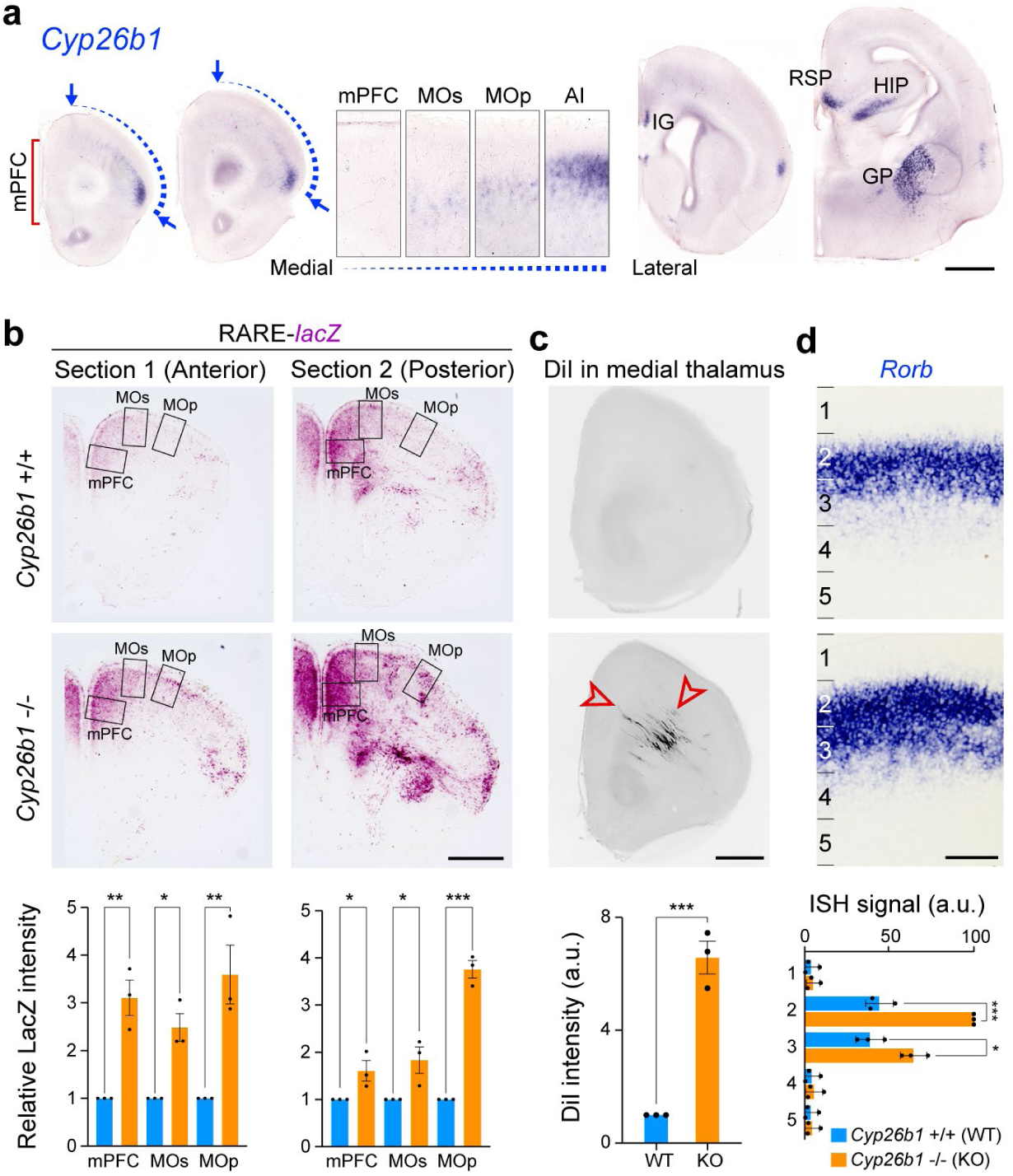
Increased retinoic acid signaling promotes mPFC-medial thalamic connectivity. **a**, Expression of *Cyp26b1* in the P0 mouse brain using *in situ* hybridization. *Cyp26b1* expression displays a gradient from the insula (AI) to MOs (insets). Arrows indicate *Cyp26b1* expression at the insula and MOs. Scale bars, 200 μm. N = 4 **b**, β-Galactosidase histochemical staining of the mPFC of *Cyp26b1*+/+ (WT); *RARE-lacZ* and *Cyp26b1*-/-(KO); *RARE-lacZ* mouse brains at 18 PCD. Significant increase in signal intensity was observed in *Cyp26b1* KO brains in the mPFC, MOs, and MOp. Signal intensity was quantified in the boxed area (mPFC, MOs, and MOp). Two-tailed Student’s t-test: WT vs. *Cyp26b1* KO: ***P < 0.0005, **P < 0.005, *P < 0.05; N = 3 per genotype; Errors bars: S.E.M. **c,** DiI labeling in 18 PCD frontal cortex after DiI was placed in the MD thalamus of WT and *Cyp26b1* KO brains. A significant increase in the density of labeled ingressing thalamocortical axons was observed in the perinatal *Cyp26b1* KO frontal cortex. Two-tailed Student’s t-test: ***P *=* 0.0004; N = 3 per genotype and condition. Scale bars: 200 μm. **d,** The expression of *Rorb*, a gene expressed preferentially in layer 4 neurons, is upregulated in *Cyp26b1* KO mice. The cortex was divided into five equal regions spanning from the pia to the ventricular zone, and signal intensity in each region is quantified for each genotype. Two-tailed Student’s t-test: WT vs. *Cyp26b1* KO: ***P = 0.0002, *P = 0.01; N = 3 per genotype; Errors bars: S.E.M. IG, Indusium griseum; GP, Globus pallidus; HIP, Hippocampus; RSP, Retrosplenial cortex.

We also observed a significant enrichment of orthologous genes specifically upregulated in the human mid-fetal frontal lobe (from Fig. 1) in the list of genes that were downregulated in the dKO mice, including those linked to synapse and axon development, such as *Cbln1*, *Cbln2, Cdh8, Nrp1*, *Pcdh17, Sema3c* and *Tnc* (**Extended Data** Fig. 4e; Fig. 1d). Furthermore, downregulated genes identified exclusively in the mPFC were significantly enriched for both orthologous genes specifically upregulated in the human frontal lobe and ASD-related genes (Extended Data Fig. 4e). Finally, genes upregulated exclusively in the OFC were significantly enriched for both ASD- and neuroticism-related genes (Extended Data Fig. 4e).

Since the role of RA in proliferation has been previously investigated^24, 25, 33^, we analyzed the role of RA in synaptogenesis by quantifying synapse number in multiple regions of the mouse P0 cortex between dKO mice and WT littermate controls, and identified a significant reduction of PSD-95-positive excitatory synapses in the mPFC (27.6% reduction) and MOs (32.4% reduction), but not in the OFC, primary motor area (Mop), and primary somatosensory area (SSp) (Two-tailed Student’s t-test: WT vs. dKO: P= 0.0006 for mPFC; P = 0.002 for MOs; P = 0.09 for OFC; P= 0.7 for MOp; P= 0.9 for MOp; Fig. 3d, Extended Data Fig. 6b). Overall, these analyses suggest a specific role for RA signaling in the regulation of synaptogenesis and, possibly, axon development, particularly in the mPFC.

### *Rxrg* and *Rarb* are required for development mPFC-MD thalamus connectivity

We further investigated the role of RA signaling on long-range connections from the mPFC using Diffusion Tensor Imaging (DTI). Various tracing studies have identified connections to the contralateral mPFC, thalamus, nucleus accumbens, and basolateral amygdala as the main output from the mPFC^1–4, 21^. We identified a profound reduction in long-range connections between the mPFC and thalamus in dKO compared to WT mice at P5 (Paired t-test: WT vs. dKO: P *=* 0.001; Fig. 4b). There was no difference in connections between the left and right mPFC at P5 (Fig. 4c). Due to limitations of the technique and underdeveloped axon pathways at this age, we were unable to study the connections between the mPFC and nucleus accumbens, or basolateral amygdala using DTI. Our attempts to use DTI to investigate connections at later ages were hindered by technical air bubble artifacts that arose during preparation of older brains that we were not able to solve in time for this submission.

Thus, we then performed anterograde axon tracing experiments at P21, where a fluorescent lipophilic dye DiI was placed in the mPFC or medial thalamus (Fig. 4d,e; Extended Data Fig. 6) both to confirm the DTI finding and assess if the reduced mPFC-MD connectivity persists at older ages. When tracer was placed in the mPFC, there was a selective loss of labelled processes in two nuclei in the thalamus, likely the mediodorsal (MD) and anteromedial (AM) nucleus in dKO mice compared to controls (67% reduction, Two-tailed Student’s t-test: WT vs. dKO: P = 0.009), as well as a reduction of fibers in the internal capsule (Fig. 4d; Extended Data Fig. 6c). Placement of lipophilic tracer in the medial thalamus showed reduction of labeled processes in the ventromedial PFC, specifically infralimbic and prelimbic areas (96.2% reduction, paired t-test: WT vs. dKO: P = 0.003) with fibers present in the anterior white matter for both (Fig. 4e; Extended Data Fig. 6d). Because there is reduced RA activity in the outer shell of the striatum in the dKO brain (Extended Data Fig. 3a), we assessed the trajectories of axon fibers through the striatum. We found that fibers transit through the medial aspect of the striatum (Extended Data Fig. 6c,d), suggesting that alterations in RA signaling in the striatum do not affect guidance of reciprocal mPFC-MD connectivity. In addition, reduction of connections between the mPFC and thalamus was not due to cell death in the mPFC (Paired t-test: WT vs. dKO: P = 0.618; Extended Data Fig. 7e).

While we saw no changes in RA activity in the dKO thalamus (Extended Data Fig. 3a), RA signaling has been previously implicated in thalamic development^53^, thus we assessed whether other thalamocortical connections were altered in the dKO mice at P5 using DTI. There was no difference in thalamocortical connectivity with the MOp, primary auditory area (AUDp), or SSp (Paired *t*-test: WT vs. dKO: P *=* 0.7, 0.6, 0.3 for MOp, AUDp, and SSp, respectively) (Fig. 4a,c; Extended Data Fig. 7b). Consistent with this, formation of barrel fields in the SSp in P5 control and dKO brains showed no significant difference (Extended Data Fig. 7c). Given that *Aldh1a3* is expressed in the ACA^51^, we also examined whether thalamocortical innervation was altered in the ACA, but found no difference in the number of streamlines (Paired t-test: WT vs. dKO: P *=* 0.3) (Fig. 4a; Extended Data Fig. 7b). The corticospinal tract (CST) and connections across the corpus callosum between left and right mPFC, SSp and MOp in the dKO showed no difference in the number of streamlines compared to control (Paired t-test: WT vs. dKO: P *=* 0.5, 0.4, 0.5, 0.12 for CST, mPFC-mPFC, SSp-SSp, and MOp-MOp, respectively: Fig. 4a,c, Extended Data Fig. 7b). Furthermore, scalar indexes, which describe the microstructural integrity of white matter, were similar between WT and dKO mice in the corpus callosum, anterior commissure and internal capsule (Extended Data Fig. 7a). In addition, the width of the CST at P30 was slightly increased in the dKO mice (Two-tailed Student’s t-test: WT vs. dKO: P *=* 0.03; Extended Data Fig. 7d). In summary, deletion of *Rxrg* and *Rarb* leads to reduction of RA signaling specifically in mPFC, as well as selective reduction of reciprocal mPFC-MD connectivity.

### Ectopic retinoic acid signaling expands medial thalamic connectivity and promotes layer 4 molecular identity

In primates, MD innervation of the frontal cortex is expanded laterally compared to rodents, with the primate PFC areas have a more prominent layer 4 ^16, 18, 21^. Thus, we investigated whether expansion of RA signaling is sufficient to increase and laterally extend MD innervation and alter layer 4 development in the mouse neonatal frontal cortex by genetically deleting the RA-degrading enzyme CYP26B1 in mice (Extended Data Fig. 8a). Whereas heterozygotes appear normal and are fertile, mice with constitutive deletion of *Cyp26b1* exhibited various body malformations and died perinatally. Nevertheless, it has been previously shown that the genetic deletion of *Cyp26b1* in mice resulted in expansion of RA signaling in multiple organs prenatally^54^ and the postnatal mouse mPFC^38^. Consistent with previous findings in humans^32^ and mice^38^, *CYP26B1* is upregulated in M1C compared to the PFC during human mid-fetal development (Fig. 1g) and upregulated in the anterior insula and MOs/MOp of the neonatal mouse frontal neocortex (Fig. 5a; Extended Data Fig. 8b). To investigate the possible role of CYP26B1 in restricting RA signaling to the mouse medial frontal cortex, we generated *Cyp26b1* KO mice that also harbored the *RARE-lacZ* transgene. The lack of *Cyp26b1* resulted in spreading of RA signaling dorso-laterally toward the MOs and MOp regions of the *RARE-lacZ* reporter mouse line at 18 PCD (Two-tailed Student’s t-test: WT vs. KO: P *=* 0.004, 0.04 for mPFC in section 1 and 2, respectively; P *=* 0.006, 0.04 for MOs in section 1 and 2, respectively; P *=* 0.01, 0.0001 for MOp in section1 and 2, respectively; Fig. 5b; Extended Data Fig. 8a). This increase was not robustly observed in more posterior regions (Extended Data Fig. 8c).

Next, we used *Cyp26b1* KO mice to study the effects of expansion of RA signaling into the dorso-lateral frontal cortex, by placing DiI into the medial thalamus of fixed postmortem WT and KO brains, harvested at E18 due to perinatal lethality of the KO mice. Histological analysis of the E18 brains revealed that WT littermates had occasional thalamocortical axons within the medial and dorso-lateral frontal white matter and cortex at this age. In contrast, KO mice showed precocious and robust innervation of both the medial and dorso-lateral frontal cortex by the axons originating from the medial thalamus (657% increase; Two-tailed Student’s t-test: WT vs. KO: P *=* 0.0004; Fig. 5c, Extended Data Fig. 9b). We also observed moderately enlarged frontal cortex (Two-tailed Student’s t-test: WT vs. KO, P *=* 0.0001 for frontal cortex size) and grossly typical cytoarchitecture of the cortical wall and cortical plate analyzed areas of the *Cyp26b1* KO cortex (Extended Data Fig. 10a, b).

Furthermore, we observed an upregulation and expansion in laminar expression of layer 4 marker, *retinoic acid-related orphan receptor beta* (*Rorb*), in the frontal cortex of *Cyp26b1* KO mice (Two-tailed Student’s t-test: WT vs. KO, P *=* 0.0002 in bin 2; P *=* 0.01 in bin 3; Fig. 5d; Extended Data Fig. 9a). Similarly, misexpression of a plasmid expressing *Aldh1a3*, in the dorso-lateral fronto-parietal cortex using *in utero* electroporation lead to the expansion of *Rorb* expression domain and ectopic expression of *Rorb* in deep layers at P0 (Two-tailed Student’s t-test: control vs. *Aldh1a3*-electroporated: P *=* 0.0001; Extended Data Fig. 10c).In summary, we identified that ectopic RA signaling in the perinatal mouse frontal cortex leads to expansion of thalamocortical innervation as well as regional and laminar expansion of expression of layer 4 marker, *Rorb*,, both characteristics of the lateral granular PFC in anthropoid primates^16^.

## Conclusion

In this study, we report that spatial regulation of RA concentration and signaling during the mid-fetal period is important for PFC development. We identified a high anterior to low posterior gradient of RA in the developing primate cortex and of RA receptors, *Rxrg* and *Rarb,* in the developing neocortex of both rodents and primates. The graded expressions of transcription factors, PAX6 and NR2F1/COUP-TF1, predominantly in early cortical progenitor cells, are required for early regional patterning of the cerebral cortex^11, 13–15, 55, 56^. It is conceivable that the anterior-posterior gradient of RA is refined by interplay with the PAX6 anterior-posterior gradient and an opposing NR2F1 posterior-anterior gradient. *Pax6* expression can be induced by RA^57^ and PAX6 positively regulates *Aldh1a3* expression^50^, whereas NR2F1 has been shown to repress RA-stimulated transcription ^57^. Furthermore, in the developing retina, RA regulates FGF8 ^58^ which was previously shown to regulate the patterning of the frontal cortex in mice^10, 12^. Thus, graded RA signaling may directly interact with previously identified cortical patterning signaling mechanisms.

We showed that *Rxrg* and *Rarb* are required for proper development of mPFC-MD long-range connectivity in mouse. Interestingly, there is a ventro-lateral and dorso-lateral expansion of *ALDH1A3* expression in the mid-fetal primate neocortex, similar to the expansion of PFC-MD connectivity in primates, and a posterior shift of *Cyp26b1* expression, co-incident with posterior expansion of the primate PFC. Long-range connectivity with the thalamus has previously been shown to be integral for specification of the primary and secondary sensory cortex^11^, and was recently reported to regulate *Cyp26b1* in the mouse frontal cortex^38^. Of note, mid-fetal cortical development in human and macaque, and equivalent perinatal/neonatal development in mice, is a critical period for laminar and areal specification of neurons, onset of dendritic growth and synaptogenesis, axon pathfinding, and ingression of thalamocortical axons^28^. Taking this evidence together, we propose expansion of RA signaling is a mechanism for expansion of association areas in the primate PFC through the expansion of innervation from the MD nucleus of the thalamus. Noteworthy, while our study has not revealed how changes in RA signaling may regulate thalamocortical innervation, we did observe several genes encoding axon guidance molecules that are enriched in the neonatal frontal cortex and dysregulated in mice lacking *Rxrg* and *Rarb.* Furthermore, we also found that an increase in RA signaling in the mouse neocortex leads to the expansion of *Rorb* expression, a key marker of granular layer 4, which is a distinct feature of the laterally expanded primate PFC^16, 18, 20, 21^.

The PFC is hypothesized to have been derived from two prime entities: the ventral paleocortical moiety that evolved from the olfactory/pyriform system and the dorsal archicortical moiety that evolved from the dorsal hippocampus^58^. Interestingly, we identified the mid-fetal frontal and temporal lobe, which are adjacent to each moiety, as having the most differentially expressed genes during mid-fetal neocortical development. Noteworthy, RA signaling is involved in the development of both entitites^36, 37^. In mice, the meninges covering the hippocampus, which resides next to the temporal neocortex, express the other two RA synthesizing genes (*Aldh1a1*/*Raldh1* and *Aldh1a2*/*Raldh2*) and RA signaling plays a critical role in hippocampal neurogenesis and function^33^. RA signaling is also required for the development of the olfactory system and its long-range connections with the ventrolateral forebrain^37^. Interestingly, we also identified an enrichment of RA in the ITC of humans, but not macaques. The ITC is an association area in the temporal lobe involved in object and face recognition and exhibits unique specialization and connectivity in humans^23, 49^. Noteworthy, the expression of the serum retinol-binding protein gene is progressively restricted to the higher-order fronto-parieto-temporal association areas during the postnatal development of macaque neocortex^59^. As expansion of the prefrontal and temporal association areas have been proposed to be one of the evolutionary underpinnings of advanced cognition^17, 19, 20, 22, 23^, it will be important to explore whether RA signaling has a broader role in developmental specification and expansion of association areas.

Clinically, disruptions in PFC development and reciprocal long-range connections between the MD nucleus of the thalamus and the PFC are thought to underlie cognitive dysfunction in schizophrenia^9^ and pathophysiology of ASD^5^. Disruption of RA signaling has been implicated in schizophrenia^45–48^, including enrichment of rare variation mutations in *RARB* in cases with severe cognitive deficits^48^. Furthermore, RA is known teratogen that can cause defects in human cerebral cortical development^60^. Our findings identify a possible mechanistic link between RA dysregulation, cognitive dysfunction, and neuropsychiatric and neurodevelopmental disorders.

## Methods

### Human developmental brain RNA-seq analysis

Bulk-tissue human brain developmental RNA-seq data (counts file) with its metadata information was downloaded from development.psychencode.org. A total of 73 mRNA samples corresponding to 11 possible neocortical areas/regions from windows 3 and 4 (16-22 PCW) were considered for analyses (Extended Data Fig. 1). The human neocortical areas under study are orbital (oPFC/OFC), dorso-lateral (dlPFC/DFC), ventro-lateral (vlPFC/VFC), medial (mPFC/MFC) prefrontal cortex, and primary motor cortex (M1C) from the frontal lobe; primary somatosensory cortex (S1C) and posterior inferior parietal cortex (IPC) from the parietal lobe; primary auditory cortex (A1C), posterior superior temporal cortex (STC), and inferior temporal cortex (ITC) from the temporal lobe; and primary visual cortex (V1C) from the occipital lobe. A TMM normalization procedure was applied (function *normalizeCounts* from tweeDEseq package in R) to the expression of 15,724 protein-coding genes that show sufficiently large counts (determined with function *filterByExpr* from edgeR package in R). To identify genes that are upregulated in a given brain lobe, we first applied RNentropy^61^ available as a package in R, to determine which genes are differentially expressed among the 11 neocortical areas. Then, we considered a gene to be specifically overexpressed in a given lobe if i) there is at least one area in this lobe where the gene is significantly upregulated, ii) the gene is not upregulated in any area of the other lobes, and iii) the gene is under-expressed in at least 30% of the areas from the remaining lobes. Similarly, we identified genes that are specifically upregulated in the PFC compared to the M1C, or vice versa, by first running RNentropy pairwise comparisons between M1C and each of the prefrontal areas independently. Then, a gene was considered to be upregulated in PFC if i) it was upregulated in a prefrontal area in at least one of the comparisons, ii) it was not upregulated in M1C in any of the comparisons, and iii) it was under-expressed in M1C in at least three of the comparisons. A gene was considered to be upregulated in M1C if i) it was overexpressed in MOp in at least three of the comparisons, ii) it was not upregulated in any PFC area, and iii) it was under-expressed in a prefrontal area in at least one of the comparisons. Principal component analyses were performed using the *prcomp* function in R by centering the log2-transformed expression data of the selected genes. Significant GO terms were obtained via *goana* function from the limma package in R, reporting the ones with at least 10 genes in the background and at least 5 in the dataset. Sequencing data were deposited at http://psychencode.org and NCBI dbGAP Accession phs000755.v2.p1

### Animals

All experiments using animals were performed in accordance with protocols approved by the Yale University Institutional Animal Care and Use Committee (IACUC). The day on which a vaginal plug was observed was designated as 0.5 post conception day (PCD). *RARE-lacZ* (Tg(RARE-Hspa1b/lacZ)12Jrt) mice, and Timed pregnant CD-1 mice for *in utero* electroporation were purchased from Jackson Laboratory and Charles River Laboratories, respectively.

### Generation of *Rxrg*, *Rarb*, and *Cyp26b1* knockout mice using CRISP-Cas9 gene editing technique

The overall strategy for the generation of *Rxrg* and *Rarb* KO mice follows a previously described protocol using CRISPR-Cas9 genome editing technique^62^. For the construction of the templates of guidance RNA, two sets of top and bottom strand oligomers (see **Supplementary Table 4**) directing the double strand break at targeting sites were annealed and ligated into *BbsI* site of pX330 vector^63^. After amplification of insert with T7-tagged primers (see **Supplementary Table 5**), guidance RNAs were synthesized by T7 RNA polymerase. The coding region of Caspase9 was PCR amplified using pX330 as a template and inserted into the pSP64 Poly(A) vector (Promega). Vectors were digested and linearized with *EcoRI*. Caspase mRNA was synthesized by SP6 RNA polymerase. Guidance RNAs and Cas9 mRNA were purified by MEGAclear Transcription Clean-Up Kit (Ambion). Cas9 mRNA, and two guidance RNAs were mixed at a concentration of (10 ng; 100 ng; 100 ng μl^−1^) in the microinjection buffer (5mMTris-HCl pH7.5; 0.1M EDTA) and injected into the pronuclei of fertilized eggs from B6SJLF1/J mouse strain. The first generation (F0) mice with recombined alleles were identified by PCR with two primer sets designed outside and inside of targeted area (**Supplementary Table 5**; Extended Data Fig. 2d,9a), confirmed by sequencing. The germ line transmission in F1 generation was confirmed by the same sets of PCR primers. For generation of *Rxrg* KO mice, a pair of guidance RNAs flanking whole exon 3 and 4 were designed to delete a large part of DNA binding domain (Extended Data Fig. 2d) ^29^. For generation of *Rarb* KO mice, a pair of guidance RNAs were designed to delete the whole of exon 9 and a part of exon 10 (Extended Data Fig. 2d). As a result, α-helical sheets of H4 to H8 in the ligand binding domain were deleted and a frame shift occurred in the rest of C-terminal region, which results in total abolition of receptor activity^64^. For generation of *Cyp26b1* KO mice, a pair of guidance RNAs were designed to delete the whole of exon 3 and 6 as described previously to abolish enzymatic activity (Extended Data Fig. 8a)^65^. All primer sequences are listed in Supplementary Table 5.

### *In situ* hybridization

Section *in situ hybridization* was performed as described previously^66^. Antisense digoxigenin (DIG)-labeled RNA probes were synthesized using DIG RNA Labeling Mix (Roche). Human and mouse *ALDH1A3* (Clone ID 6208628, and 6515355, respectively), *RXRG* (Clone ID 4635470, and 5707723, respectively), *RARB* (Clone ID 30341884, and 30608242, respectively), and mouse *Rorb* (Clone ID 5358124), *Cbln2* (Clone ID 6412317), *Cyp26b1* (Clone ID 6400154) cDNAs were purchased from GE Healthcare. Mouse *Meis2* DNA was a gift from John Rubenstein. For macaque *in situ* hybridization, human probes were used because of high similarity between human and macaque transcripts (95.4% identity in *ALDH1A3*; 97.7% in *RXRG*; 98.8% in *RARB*). Sections were obtained from 21, 22 PCW human brains and 110, 114 PCD macaque brains. *In situ* hybridization were repeated using these 2 sets. *Rorb* intensity was quantified using Image J. *Rorb* thickness was quantified by dividing the cortical plate into 5 equal bins and intensity in each bin was quantified using Aperio ImageScope (Leica).

### Enzyme-linked immunosorbent assay (ELISA)

Eleven neocortical areas were dissected from four fresh frozen postmortem human mid-fetal brains (16,18, 18 and 19 PCW) and four fresh frozen macaque brains (all four 110 PCD) as described in Zhu et al. 2018^67^. Each sample was further microdissected into three pieces and weighed. Each piece was independently homogenized using a dounce homogenizer in three to four volumes of homogenizing buffer (isopropanol: ethanol=2:1; 1 mg/ml butylated hydroxytoluene), followed by centrifugation at 10K rpm for 10 minutes at 4 °C. Supernatant was used for both determination of protein concentration by BCA kit (Thermo-Fisher Scientific), and RA concentrations using ELISA colorimetric detection Kit according to the manufacturer’s instructions (Cat. MBS705877, MyBioSource, San Diego, CA). This kit could not distinguish forms (all trans retinol, all trans retinal, and all trans retinoic acid). Thus, the concentration of RA was the overall concentration.

### Quantitative reverse transcription-PCR

Total RNA was isolated from freshly microdissected cortices after removal of the olfactory bulb and striatum using Trizol (Thermo-Fisher Scientific). cDNAs were prepared using SuperScript II (Invitrogen) from three independent WT cerebral hemispheres. Quantitative reverse transcription (RT)-PCR was performed as described previously^68^. At least three replicates per transcript were used for every reaction. The copy number of transcripts was normalized against the house keeping TATA-binding protein (TBP) transcript level. For *Rxra,b,g, Rara,b,c* and *Tbp* primer sets, correlation (R2) was higher than 0.98, and the slope was −3.1 to −3.6 in each standard curve. Primers to detect the expression of the genes above were designed in a single exon. Primer sequences are listed in **Supplementary Table 5**.

### β-Galactosidase histochemical staining

Brains were dissected from P0 *RARE-lacZ* mouse pups and drop-fixed in 4% paraformaldehyde for 2 hours at 4C, followed by embedding in OCT. Brains were sectioned at 20 µm by cryostat (Leica CM3050S) after they were frozen. β-Galactosidase staining followed the protocol described by Kokubu et al^69^. We used Red-gal (Sigma-Aldrich) for the chromogenic reaction. Intensity of β-Galactosidase staining was quantified using Image J.

### Nissl staining

Postmortem brains were harvested and fixed with 4% paraformaldehyde overnight at 4°C, followed by embedding in OCT. Brains were sectioned at 15-20 µm by cryostat (Leica CM3050S) after they were frozen. After PBS wash, sections were dehydrated using increasing concentration of ethanol, followed by cresyl violet, wash, and second ethanol dehydration.

### Immunohistochemistry

The brains were dissected from each embryo and fixed with 4% paraformaldehyde overnight at 4°C, followed by embedding in OCT. Brains were sectioned at 15-20 µm by cryostat (Leica CM3050S) after they were frozen.

The sources of primary antibodies were anti-L1CAM (1:500; Millipore), anti-cleaved capase3 (1:500; Cell Signaling), anti-PSD95/DLG4 (1:500; Invitrogen). Secondary antibodies: Alexa Fluor 488- or 594-conjugated AffiniPure Donkey anti-Rabbit IgG (Jackson ImmunoResearch). For all microscopic analysis, LSM510 META (Zeiss), and LSM software ZEN were used.

### Quantification of postsynaptic puncta marked by PSD-95/DLG4 immunostaining

For each region of both WT and dKO mice, using the 488 nm channel to detect DLG4/PSD-95 and DAPI to label nuclei, seven serial optical sections at 0.8 μm intervals over a total depth of 5 μm were imaged and the 2^nd^, 4^th^, and 6^th^ images were eliminated from further analysis to avoid overlap in counting^70^. Area of each image is 0.079 mm^2^. The number of PSD-95 puncta on each image was counted automatically using ImageJ using threshold of 985 to 4095 and analyze particles function. At least two sections from each animal were selected for counting, and at least 3 animals for each genotype were used.

### Anterograde tracing of axons

For anterograde tracing of axons between mPFC and thalamus, brains were collected at either 18 PCD or P21, and fixed overnight in 4% paraformaldehyde at 4 °C. Brains were then hemidissected. A crystal of 1,1#-dioctadecyl-3,3,3#,3# tetra-methyl-indocarbocyanine perchlorate (DiI, Sigma-Aldrich) was inserted either into the mPFC, MD nucleus of the thalamus, or medial thalamus under the stereomicroscope. The size of the crystal is ∼200 μm. Brains were then placed in 1% paraformaldehyde in PBS and left for 14 days at 37 °C. Following DiI diffusion, the brains were sectioned coronally on a vibrating microtome (Leica) at 80 μm thickness and stained with DAPI. Sections were mounted onto glass and immediately sealed in VECTASHIELD Hardset Antifade Mounting Medium (VECTOR Laboratories). Slides were analyzed under ApoTome.2 microscope (Zeiss) and intensity was quantified using ImageJ.

### Plasmid construction

For construction of expression vectors, full-length cDNAs (mouse *Aldh1a3*, Clone ID 6515355, purchased from GE Healthcare) were inserted into pCAGIG vector (pCAGIG was obtained from Addgene (Plasmid # 11159).

### *In utero* electroporation

*In utero* electroporation was performed as previously described^68^. Plasmid DNA (4 μg/μl) was injected into the lateral ventricle of embryonic mice at E13.5–E14.5 and transferred into the cells of the ventricular zone by electroporation (five 50-ms pulses of 40 V at 950-ms intervals). Brains were dissected at P0. Brains and tissue sections of electroporated animals were analyzed for GFP expression after fixation with 4% paraformaldehyde at P0.

### Mouse RNA-seq data generation and analysis

Mouse brains were dissected at P0 in ice-cold sterile PBS, fresh frozen, and stored at −80 °C. Brains were incubated in RNAlater-Ice at −20 °C for 12-16 hours prior to further dissection. mPFC, MOs, and OFC were microdissected based on Paxinos and Frankin, 2007^71^ and the Allen Mouse Brain Atlas (mouse.brain-map.org/static/atlas)^72^ and RNA was isolated using RNeasy Plus Micro kit with additional on-column DNAase step (Qiagen). RNA quality and amount were quantified using High Sensitivity RNA Screen Tape assay (Agilent), and concentration was standardized to 10 ng/ul. SMART-seq v4 Ultra Low Input Kit (Takara) was used to create cDNA, and concentration was quantified using Quant-iT Picogreen kit (Thermo-Fisher Scientific). Nextera XT DNA library Prep Kit (Illumina) were used to create cDNA libraries for sequencing. Libraries were normalized and sequenced at the Yale Center for Genomic Analysis (YGCA) using the NovaSeq with 100 bp paired end reads. Reads from each library were mapped against the mouse assembly GRCm38 using STAR v.2.6.0a (gtf and fasta files downloaded from Ensembl version 94; parameters: --readFilesIn $j1 $j2 -- outSAMattributes All --outFilterMultimapNmax 1 --outSAMstrandField intronMotif -- outFilterIntronMotifs RemoveNoncanonical --quantMode TranscriptomeSAM -- outFilterMismatchNoverLmax 0.1 --alignSJoverhangMin 8 --alignSJDBoverhangMin 1 --outSAMunmapped Within --outFilterType BySJout). Counts were obtained using featureCounts v1.6.2 with -p parameter.

To compare the gene expression patterns of three WT vs three *Rarb*/*Rxrg* dKO mice, a TMM a procedure was applied (function *normalizeCounts* from tweeDEseq package in R) to the expression of 15085 protein-coding genes that show sufficiently large counts (determined with function *filterByExpr* from edgeR package in R). We assessed DEx genes in each brain region (mPFC, OFC, and MOs) running RNentropy independently among WT and dKO mice per region. Genes overexpressed in a given condition are those that are both significantly upregulated in that condition and significantly downregulated in the opposite condition according to RNentropy. The same criterion was applied for identification of downregulated genes. Genes with an inconsistent pattern of expression between regions were excluded. Principal component analyses were performed using the *prcomp* function in R by centering the log2-transformed expression data of the selected genes. Significant GO terms were obtained via *goana* function from the limma package in R and plotted using function *GOBubble* from GOplot package in R. Fisher test enrichments calculated for RA related genes (RA synthesis: *Rdh10*, *Rdh5*, *Raldh1*, *Raldh2*, *Raldh3*, *Adh1*, *Adh5*, *Adh7*; RA degrading: *Cyp26a1*, *Cyp26b1*, *Cyp26c1*: RA receptor: *Rara*, *Rarb*, *Rarg*, Rxra, *Rxrb*, *Rxrg*; RA binding: *Ttr*, *Rlbp1*, *Rbp1*, *Rbp2*, *Rbp3*, *Rbp4*, *Fabp5*), genes overexpressed in individual lobes of the mid-fetal human cortex based on Fig. 1, and neuropsychiatric disease related genes (downloaded from Li et al. ^27^) in up- and downregulated genes. Genes associated with the GO terms: “axon guidance”, “axon guidance receptor”, “axon development”, and “ephrin” were manually screened for anterior to posterior gradient using developingmouse.brain-map.org^73^, gensat.org^74^ and Elsen et al.^75^

### Retinoic acid receptor binding site analysis

We identified all those protein coding genes associated to GO terms related to axon guidance and development, and synapse formation among those genes upregulated in the mid-fetal human frontal cortex or up- and down-regulated in the *Rxrg* and *Rarb* dKO mice, whose human orthologs were identified using Ensemble Biomart (∼430 genes). This produced six lists of genes of which we then collected H3K27ac peaks active in mid-fetal human dlPFC/DFC from a collapsed PsychENCODE Chip-Seq dataset reported in Li et al., 2018^27^ containing dorsal frontal cortex and cerebellum samples from mid-fetal and adult brains. From the hg38 coordinates of each of the ∼1700 peaks obtained, we downloaded their DNA sequence from the UCSC Genome Browser using *twoBitToFa*^76^. We then run FIMO with default parameters to predict TFBS in those sequences using 8 JASPAR motifs associated to retinoic acid receptors in meme format^77^. For each gene and motif, we counted how many matches we observed in its putative *cis*-regulatory elements (CRE). We divided the number of bases covered by a given motif by twice the total length of each gene CREs, because our search space was in both the forward and reverse strand. We then discretized the proportion of sites spanning putative RARB/RXRG binding sites in five quantiles by including the union of the gene lists and calculated the proportion of genes falling in the five quantiles, plus a category representing genes with no predicted binding sites, and a category representing genes with no identified CREs. To compare these proportions with a null expectation, we performed randomization among the ∼15,000 protein coding genes sufficiently expressed in human frontal cortex and used in the differential expression analysis. The randomization strategy consisted of i) sampling 100 times a number of genes equal to the union of the six gene lists and, ii) for each of those random groups, sample a number of genes equal to the size of each individual list. This strategy could better approximate the notable gene overlap observed among those gene sets. Finally, we classified each gene from the randomized groups in the same quantiles categories derived from the union of the six gene lists. For each motif, we analyzed which gene lists presented categories with proportions falling outside the 95% confidence interval of the 100 randomizations. We tested for enrichment of RARB/RXRG binding sites around genes (−500 bp/+100 bp and +/-10,000 bp of the transcriptional start site (TSS)) using RcisTarget^78^, and in the H3K27ac peaks from the previous section with HOMER findMotifsGenome.pl^79^.

### Diffusion-weighted magnetic resonance imaging and tractography

Following proper euthanasia, five dKO homozygotes and five WT C57BL/6 mice at P5 were sacrificed and brains were dissected. Brains were drop fixed in a paraformaldehyde solution of 4% in 0.1 M phosphate buffered saline (PBS) for 48 hours. They were subsequently transferred to 0.1 M PBS and just before imaging to Fomblin (Sigma-Aldrich St Louis, MO, USA). The diffusion-weighted images were acquired on a Bruker BioSpin 9.4 T MRI (Bruker GmbH, Ettlingen, Germany) using a standard 3D Stejskal-Tanner spin-echo sequence with 30 different angles of diffusion sensitization at b value of 1000 s/mm^2^ and the following parameters: repetition time=2000ms; echo time=25.616ms; diffusion encoding duration = 4ms. The in-plane resolution was 0.11mm and slice thickness was 0.22mm. Overall scanning time was ∼24hours.

### Image processing and tractography

Cerebral cortical regions of interest (ROI) and thalamus were manually defined according to Paxinos and Frankin^71^ and the Allen Mouse Brain Atlas (mouse.brain-map.org/static/atlas) by D.A. and K.P. without prior knowledge of the experimental groups. The reconstruction of axonal pathways was performed with MRtrix3^80^ software using constrained spherical deconvolution^81^ and probabilistic tracking (iFOD2) with FOD amplitude cut-off of 0.1. The thalamus was used as a seeding point and each cortical ROI was used as a termination mask. To evaluate the integrity of the major white matter tracts between the groups, both internal capsules, anterior commissure and corpus callosum were manually delineated according to Paxinos and Frankin, 2007^71^ and the Allen Mouse Brain Atlas (mouse.brain-map.org/static/atlas)^72^ by D.A. and K.P. without prior knowledge of the experimental groups. Values of the fractional anisotropy (FA), apparent diffusion coefficient (ADC), radial (RD) and axial (AD) diffusivity were calculated using underlying scalar maps derived by MRtrix3.

### Processing, analysis, and image visualization

To allow robust visualization and analysis, images depicting DiI tracing or immunohistochemistry using antibody against PSD95 have been inverted and/or pseudo colored, as in Fig. 4, 5.

## Acknowledgements

We thank Suxia Bai and Timothy Nottoli for technical help in generation of gene-edited mouse lines; Fahmeed Hyder and Jelena Mihailovic for their assistance with MRI diffusion-weighted sequence design and conducting the MRIs; Alvaro Duque for use of equipment from MacBrainResource (MH113257); John Rubenstein for sharing reagents; and the members of Sestan laboratory for comments. This work was supported by the National Institutes of Health (MH106874, MH106934, MH110926, MH116488), and Simons Foundation (N.S.). The project that gave rise to these results received the support of a fellowship from “la Caixa” Foundation (ID 100010434). The fellowship code is LCF/BQ/PI19/11690010. Additional support was provided by the Kavli Foundation and the James S. McDonnell Foundation (N.S.).

## Author Contributions

M.S., K.P., and N.S. designed the research; M.S. and K.P. performed mouse experiments, analyzed the data; B.L.G. analyzed human and mouse transcriptomic datasets; G.S. analyzed the enrichment of binding sites for RA receptors; M.S. generated construct for mutant mice lines; X.X., performed pronuclear injection; D.A. analyzed mouse imaging data; A.M.M.S. dissected postmortem human and macaque tissues used for analysis; N.S. conceived the study; M.S., K.P., and N.S. wrote the manuscript. All authors discussed the results and implications and commented on the manuscript at all stages.

---

**Extended Data Figure 1.**
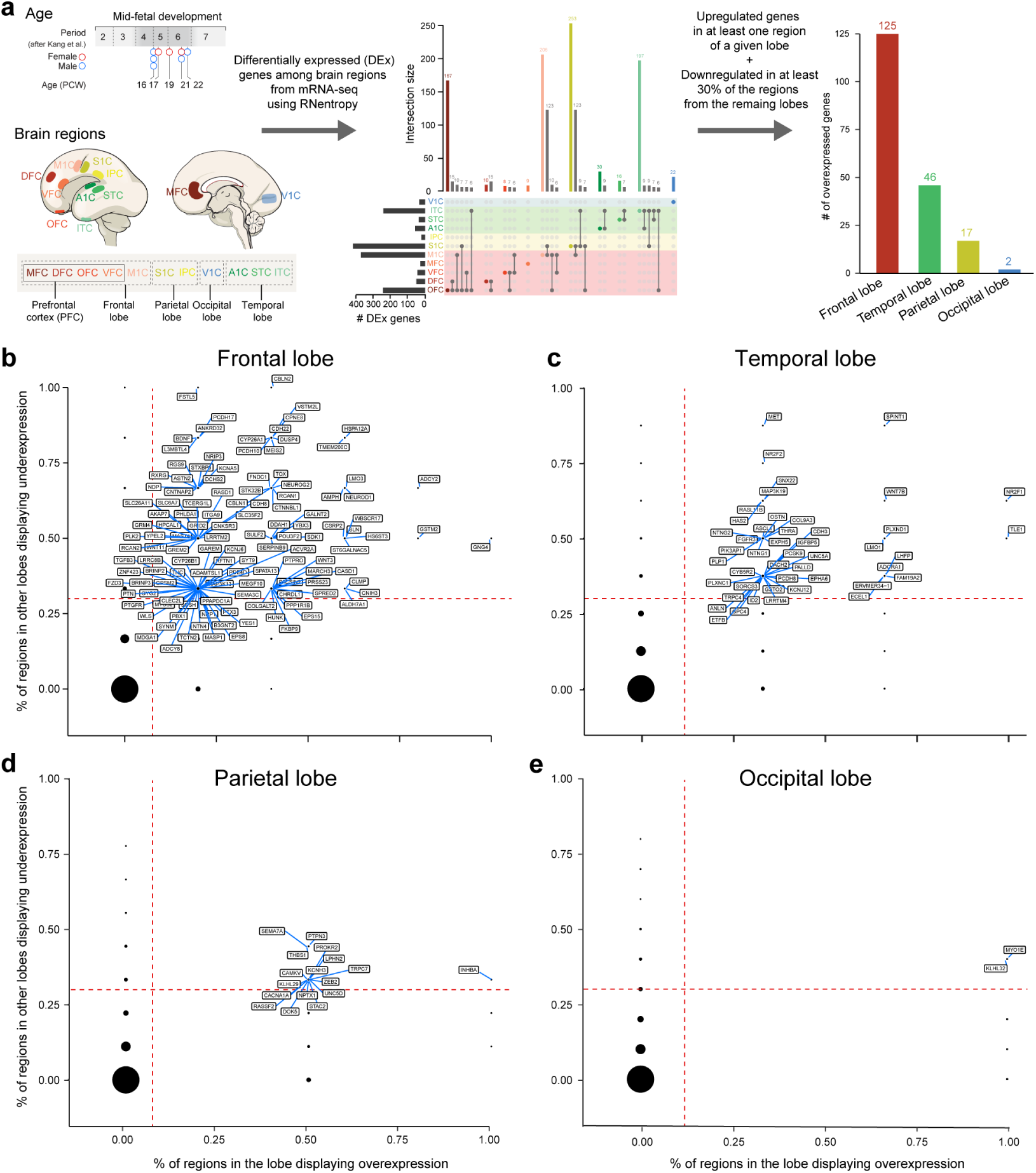
Workflow for analysis of mid-fetal human transcriptome data and genes upregulated in individual lobes. **a,** Human developmental dataset and workflow for analysis to identify genes upregulated in each cortical lobe. **b,** Genes upregulated in the frontal lobe in comparison to the other lobes. The X-axis represents proportion of regions in the frontal lobe in which the gene is significantly upregulated according to RNentropy. The Y-axis represents proportion of regions in the other lobes in which the gene is significantly downregulated according to RNentropy. Upregulated genes, the ones delimited by orange dashed lines, are labeled. **c, d, e,** Same for temporal, parietal and occipital lobes, respectively.

**Extended Data Figure 2.**
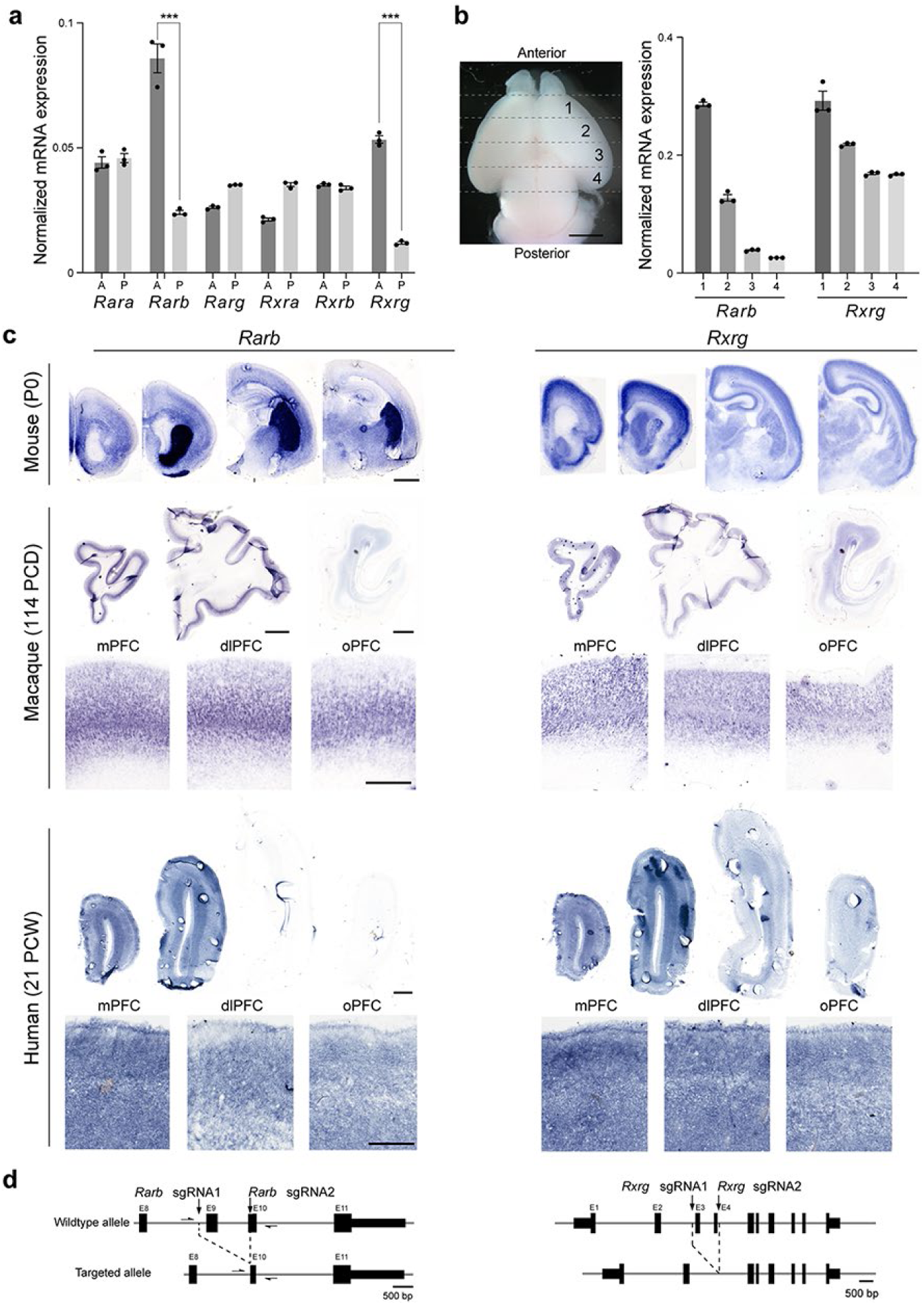
*RXRB* and *RARB* are expressed in an anterior to posterior gradient. **a,** Quantitative PCR analysis of *Rara, b, g* and *Rxra, b, g* transcripts in the anterior and posterior half of mouse cortex at P0. Note that only expression of *Rarb* and *Rxrg* showed significant enrichment in the anterior cortex **b**, Quantitative PCR analysis of *Rxrg* and *Rarb* transcripts in four sections dissected out of the cortical plate in anterior-posterior direction. Both genes showed an anterior-posterior gradient in expression level. Two-tailed Student’s t-test: ***P < 0.0005; N = 3 per condition; Errors bars: S.E.M. **c,** Expression of *Rxrg* and *Rarb* in mouse (P0), macaque (114 PCD) and human (21 PCW) brains by *in situ* hybridization. Higher magnification images of the regions of anterior cortex. *Rxrg* and *Rarb* transcripts are upregulated in the anterior part of the cortex in all three. Scale bars, 200 μm (mouse); 2 mm (human); 500 μm (human, higher magnification). N = 2 for human and macaque, N = 3 for mouse. **d**, Strategies for the generation of *Rxrg*, and *Rarb* KO mice using CRISPR-Cas9 technique^62^.

**Extended Data Figure 3.**
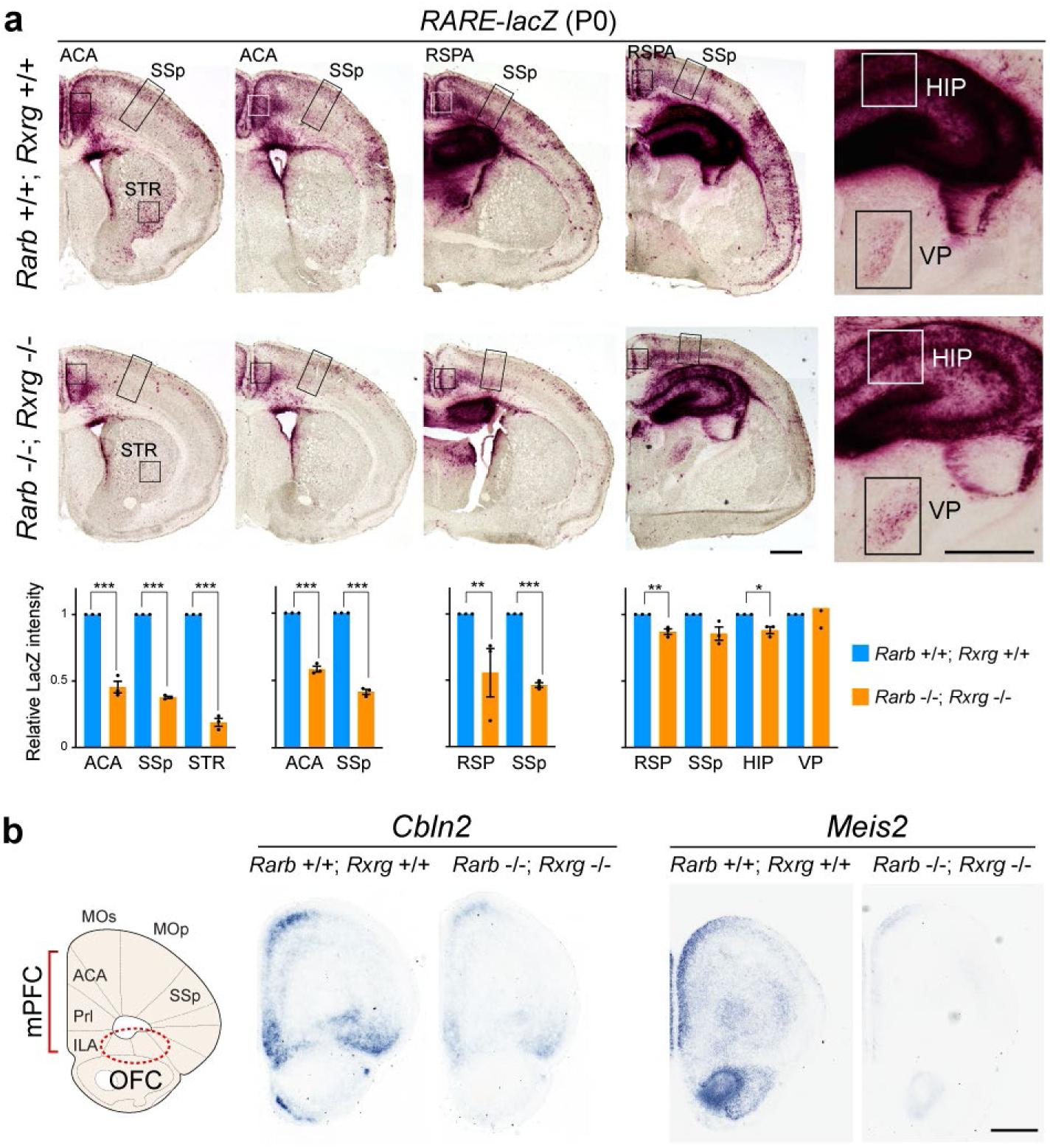
Retinoic acid signaling in posterior regions of the P0 mouse forebrain. **a**, β-Galactosidase histochemical staining of more posterior regions of *Rxrg+/+*; *Rarb+/+* (WT); *RARE-lacZ* and *Rxrg-/-*; *Rarb-/-* (dKO); *RARE-lacZ* mouse brains at P0. Signal intensity in the boxed area (ACA, SSp, RSPA, STR, HIP and VP) was quantified. Note the reduced activity in anteromedial structures including ACA and RSPA (RSP). There is also reduced expression in hippocampus and lateral striatum, but not the thalamus. Two-tailed Student’s t-test; *P = 0.009, **P < 0.005, ***P < 0.0005; N = 3 per genotype: Errors bars: S.E.M.; Scale bars, 200 μm. **b,** *Cbln2* and *Meis2* expression in P0 WT and dKO mutant brain by *in situ* hybridization at P0. Note that *Cbln2* and *Meis2* expression in mPFC was decreased in dKO. Scale bar, 200 μm. N = 3 per genotype. ACA, anterior cingulate area; HIP, hippocampus; RSPA/RSP, retrosplenial area; STR, striatum; VP, ventroposterior thalamus.

**Extended Data Figure 4.**
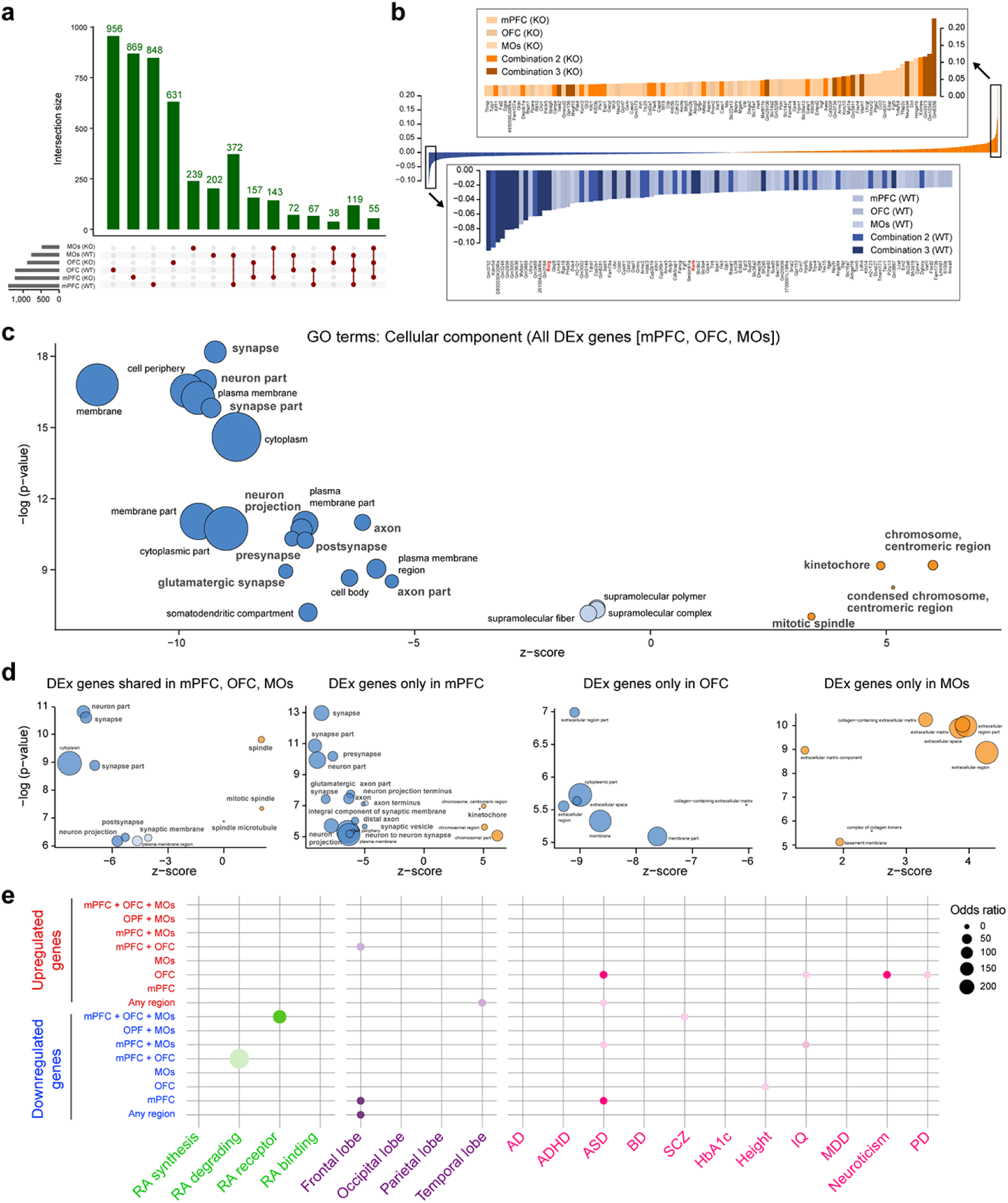
Additional analysis of RNA-seq experiments. **a**, Number of upregulated genes between P0 WT and dKO mice per region and phenotype, as well as combinations of regions and phenotypes. **b,** Gene loadings of the first principal component from PCA in Fig. 3. Colors represent the frontal cortex region where the gene was found to be upregulated. **c**, Cellular component GO terms associated with the total list of 4,768 DEx genes found, showing their z-score and nominal P values. Z-score represents the proportion of upregulated versus downregulated genes in the list of DEx genes associated to each GO term (i.e. z-score = (#up - #down) / sqrt (#all DEx associated to the GO term)). Dark blue: z-score <-5; light blue: z-score (−5,0]; orange: z-score >0. Size of the bubbles are proportional to the total number of DEx genes associated to the given GO term. **d**, Cellular component GO terms associated with DEx genes found in all three frontal cortex regions, and DEx genes unique to each region (mPFC, OFC, MOs) and their nominal P value. **e,** Enrichment of RA related genes (green), genes upregulated in individual lobes of the mid-fetal human cortex based on Fig. 1 (purple), and psychiatric disease related genes in up- and downregulated genes (pink). DEx genes are separated by genes that are DEx only in the given region (mPFC, OFC, MOs), genes that are DEx in the two given regions (mPFC + OFC, mPFC + MOs, OFC + MOs), genes that are DEx in all three regions (mPFC + OFC + MOs). Circles plotted for significant enrichments (P value <0.05), in darker color, significance is considering the adjusted P value. Diameter of circle is associated with odds ratio per legend.

**Extended Data Figure 5.**
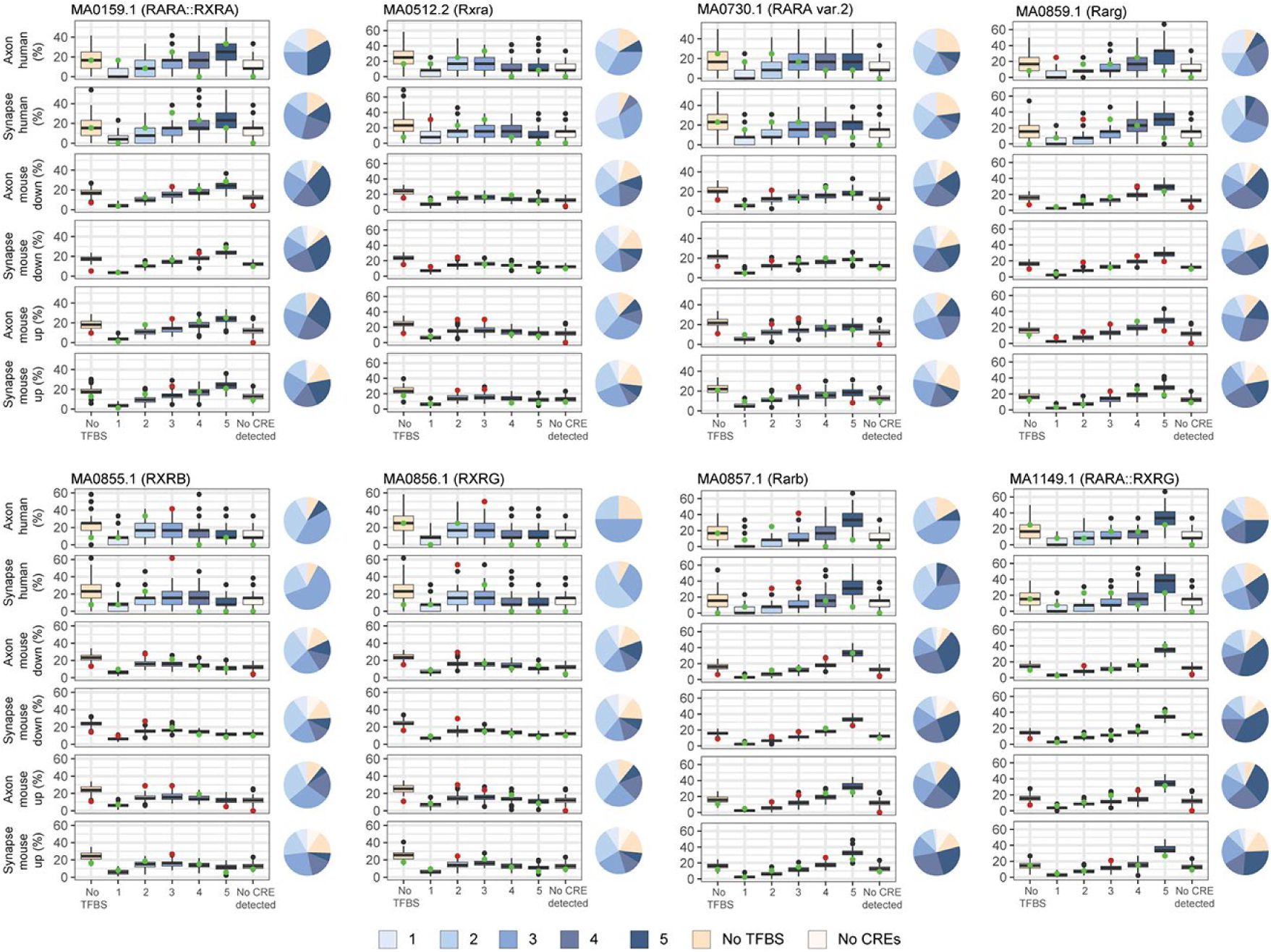
Analysis of retinoic acid receptor binding sites in regulatory regions of upregulated synaptic and axon development genes. Boxplot showing the distribution of the proportion of genes in each gene set in categories representing different levels of sequence matches for each of 8 JASPAR RA receptors binding motifs. One to five represents distribution quantiles (0.2, 0.4, 0.6, 0.8, 1) of the fraction of bases covered by RA binding sites in the union of the six gene lists. Category “No cis-regulatory element (CRE) detected” refers to genes with which we could not associate any human dlPFC/DFC mid-fetal H3K27ac peak. Colored dots represent the observed value for each category; red and green indicates whether they fall outside or inside the 95% confidence interval, respectively. Pie charts represent the observed proportion of genes in each category for each gene list and motif.

**Extended Data Figure 6.**
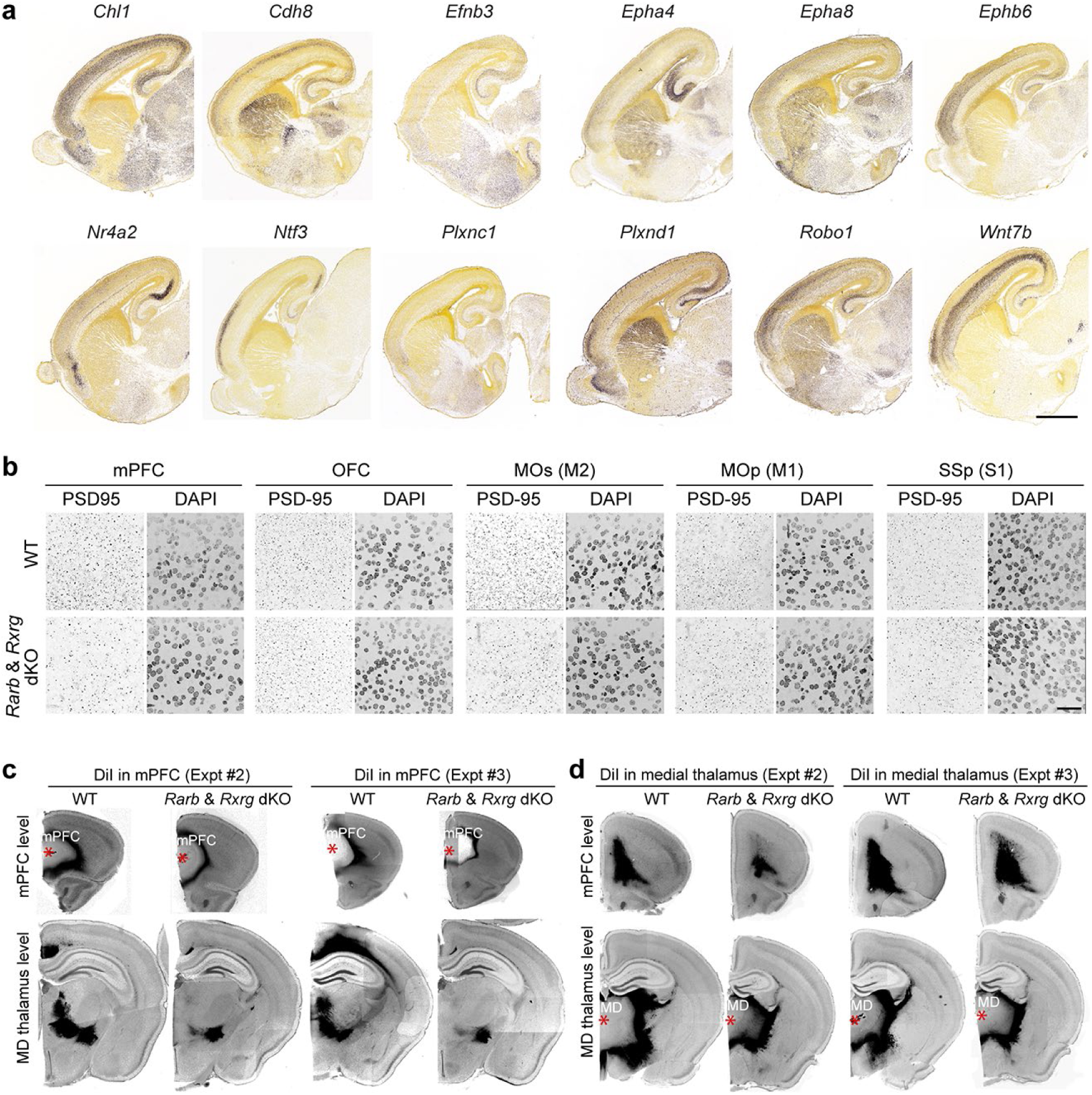
Altered synaptic density and axonal projections in *Rxrg*/*Rarb* dKO mice. **a**, Example of downregulated genes between P0 *Rxrg*+/+; *Rarb*+/+ (WT) and *Rxrg*-/-; *Rarb*-/-(dKO) that displayed an anterior to posterior gradient in 18 PCD mouse embryo (images are from the Allen Developing Mouse Brain Atlas; developingmouse.brain-map.org/). Scale bar: 1 mm. **b**, Immunostaining for PSD-95/DLG4 in the cortical subregions (mPFC, MOs, OFC, MOp, and SSp) of WT and dKO brain at P0. Each region as shown in Figure 2c. Scale bar: 25 μm. N = 3 per genotype. **c,d,** DiI placement in mPFC (**c**) and medial thalamus (**d**) with tracing data in WT and dKO brain at P21. Additional two replicates of experiment shown in Figure 4d,e are shown. Asterisks: DiI crystal placement. Scale bar: 1 mm.

**Extended Data Figure 7.**
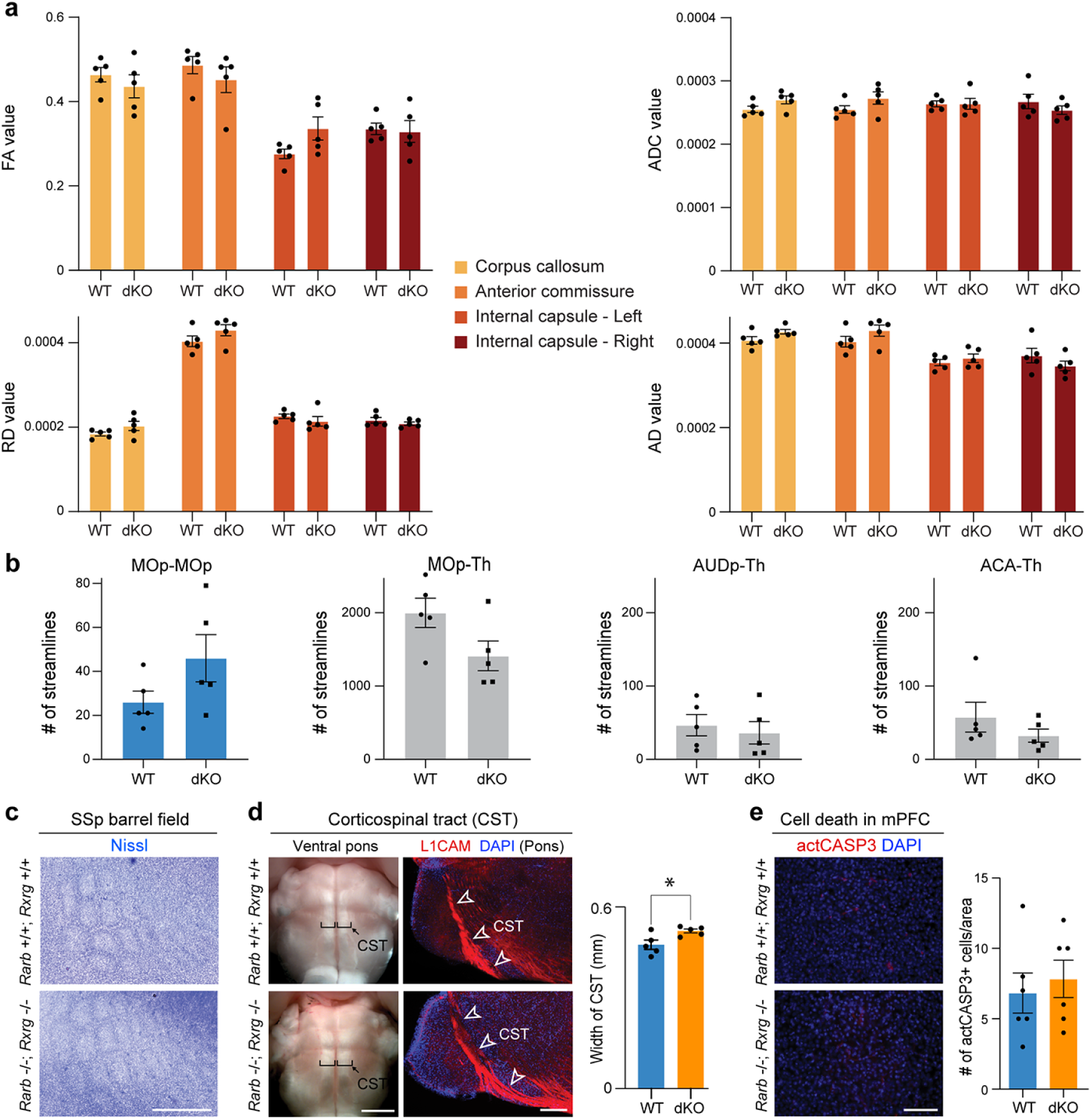
Analysis of axonal projections and cell death in *Rxrg*/*Rarb* dKO mice. **a,** Four scalar indexes which describe microstructural integrity do not differ in the four major white-matter tracts (corpus callosum, anterior commissure, left and right internal capsules) between WT and dKO mice. **b,** Number of streamlines of MOp-thalamus and corticocortical tracts, did not differ between WT and dKO. Paired t-test: NS; N = 5 per genotype; Errors bars: S.E.M. **c**, Barrel field formation was grossly normal in dKO. Barrel formation was examined by Nissl staining at P5. N = 3 per genotype. **d,** Corticospinal tract (CST) appearance at P30 was grossly normal in dKO. The width of the corticospinal tract (shown in brackets) is slightly increased in dKO. Sagittal section at P5 was stained with anti-L1CAM antibody to show the grossly normal CST formation in dKO. Two-tailed Student’s t-test: *P = 0.03; N = 5 per genotype; Errors bars: S.E.M. **e**, Frequency of apoptosis in the mPFC detected by cleaved caspase3 (actCASP3) between WT and dKO is not significantly different. Two-tailed Student’s t-test: WT vs. dKO; NS; N = 5 per genotype; Errors bars: S.E.M.; Scale bars: 500 μ m; 200 μ m (b); 1 mm (c); 100 μm (d).

**Extended Data Figure 8.**
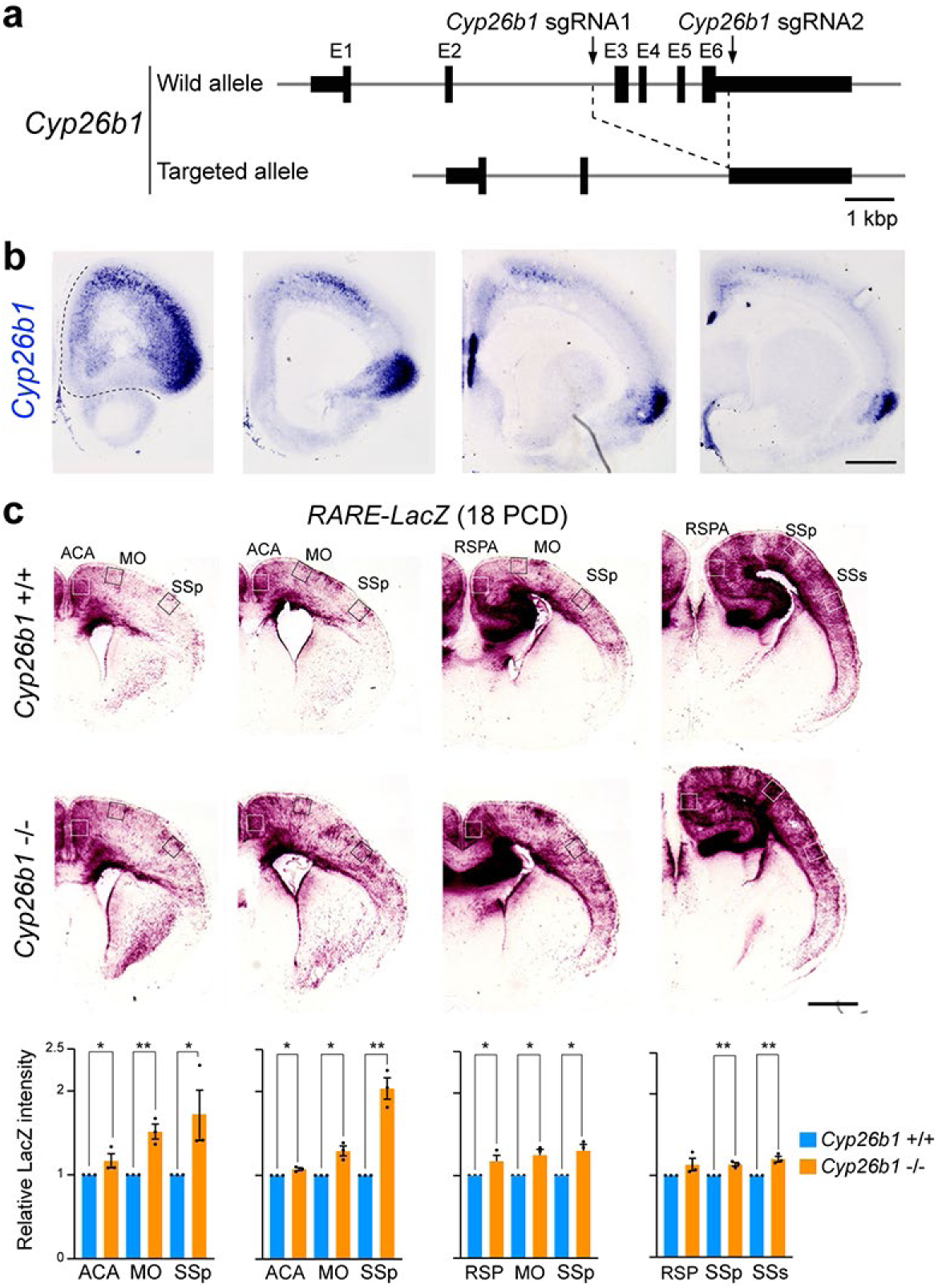
Retinoic acid signal in posterior cortical regions of *Cyp26b1* KO mice. **a**, Strategy for the generation of *Cyp26b1*-/-(*Cyp26b1* KO) mice using CRISPR-Cas9 gene editing technique^62^. **b,** *Cyp26b1* expression in P0 mice cortex by *in situ* hybridization. The colorimetric staining was purposefully extended compared to the experiment in Fig. 5a to better visualize low expressing locations. **c,** β-Galactosidase staining of more posterior regions of control *Cyp26b1*+/+; *RARE-lacZ* (Ctrl) and *Cyp26b1*-/-; *RARE-lacZ* (KO) mouse brains at 18 PCD. Scale bar: 500 μm. Intensity of signal in the boxed areas (ACA, MO, SSp) was quantified. Increase in RA signaling in *Cyp26b1* KO brains is less significant in posterior regions. Two-tailed Student’s t-test: Ctrl vs. *Cyp26b1* KO: *P < 0.05, **P < 0.005, N = 3 per genotype; Errors bars: S.E.M.; Scale bars: 200 μm.

**Extended Data Figure 9.**
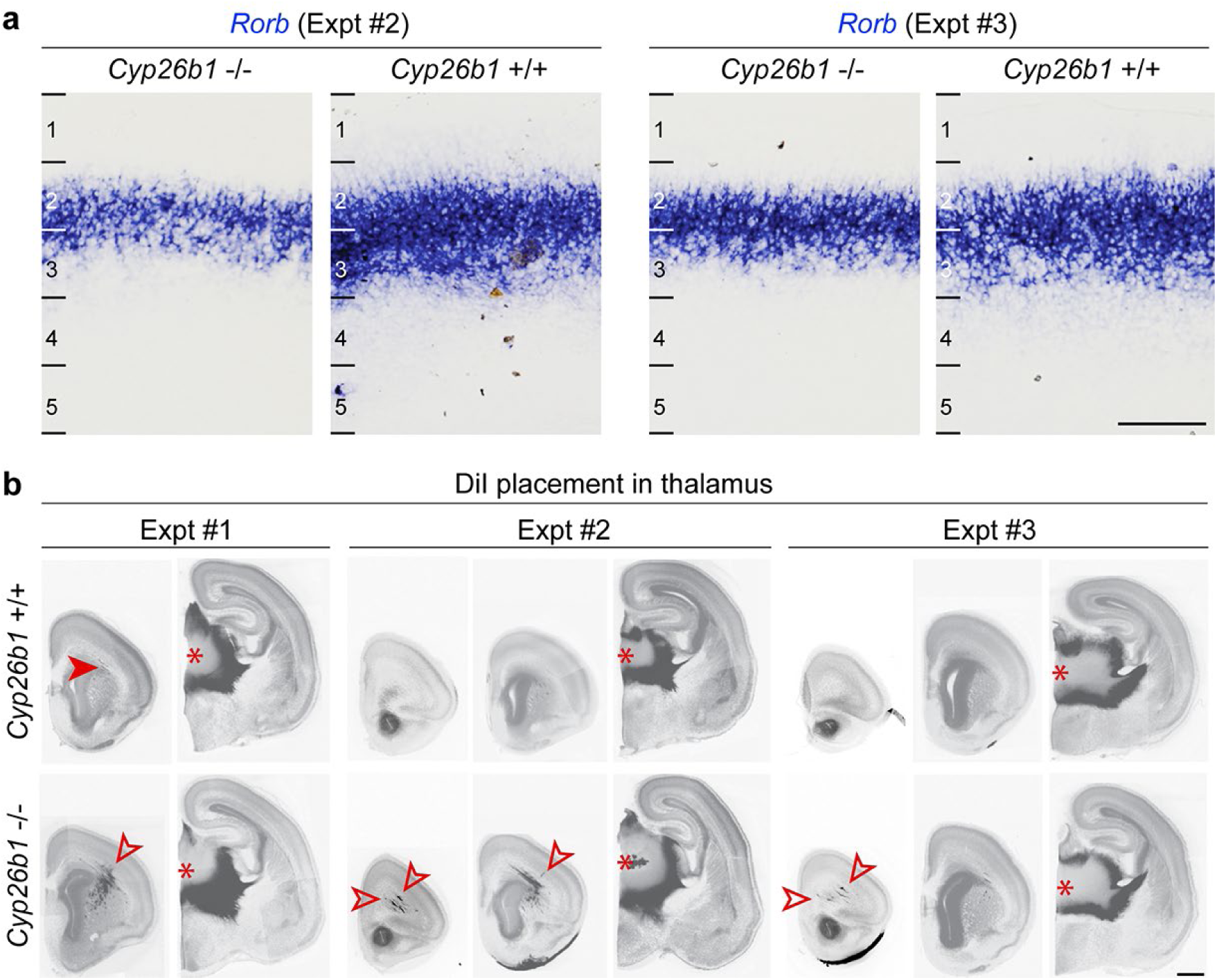
*Rorb* expression and thalamocortical projections in the perinatal *Cyp26b1* KO mice. **a,** 18 PCD, *Cyp26b1* KO mouse brains show upregulation of *Rorb*. Additional two replicates are shown. Scale bar: 100 μm. **b**, DiI was placed in the medial thalamus of WT and *Cyp26b1* KO brains, and signal was detected in the PFC. Additional two replicates of experiment in Figure 5c are shown. N = 3 per genotype and condition. Arrowheads: Thalamocortical innervation of the medial and dorso-lateral frontal cortex. Asterisks: DiI crystals placed. Scale bar: 400 μm.

**Extended Data Figure 10.**
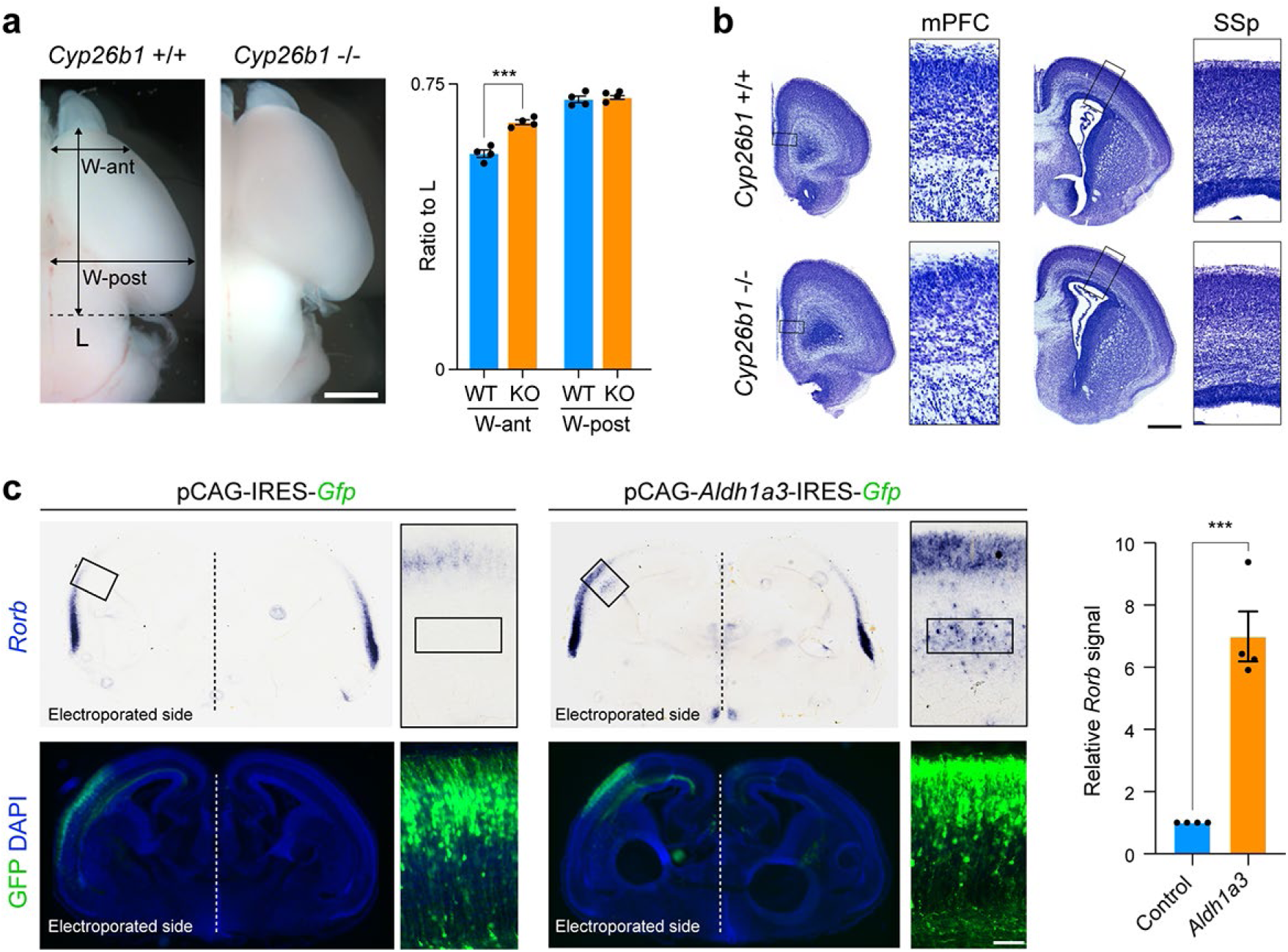
Ectopic RA signaling leads to enlargement of frontal cortex and expanded *Rorb* expression. **a**, *Cyp26b1* KO brain at 18 PCD showed enlargement of anterior/frontal cortex and not posterior cortex. Ratio of brain width at anterior cortex to total brain length was significantly increased in *Cyp26b1* KO brain compared WT. Two-tailed Student’s t-test: WT vs. *Cyp26b1* KO: ***P = 0.0001; N = 4 per genotype; Errors bars: S.E.M. **b**, Nissl staining reveals that the cortical wall and cortical plate are grossly normal when analyzed in the mPFC and SSp of *Cyp26b1* KO. **c,** Electroporation of either control pCAG-IRES-*Gfp* or pCAG-*Aldh1a3*-IRES-*Gfp* expression vector plasmid in the dorso-lateral fronto-parietal wall at 14 PCD. Brains were dissected out at P5. Misexpression of *Aldh1a3* leads to regional expansion and ectopic laminar expression of *Rorb*, when assessed by *in situ* hybridization. GFP expression as a marker of misexpressing cells are shown in lower panels. *Rorb* signal intensity in the boxed area in the cortex was quantified. Two-tailed Student’s t-test: ***P = 0.0001; N = 4 per genotype; Errors bars: S.E.M.; Scale bars: 500 μm (a); 200 μm (b); 1 mm (c); 40 μm and 500 μm (d).

